# Chromatin- and actin-mediated mitochondrial streaming leads to patterning of mitochondrial distribution in oocytes

**DOI:** 10.64898/2026.02.11.704717

**Authors:** In-Won Lee, Morteza Nazari, Jazmine Yuson, Sanjeev Uthishtran, Ross Stocker, Deepak Adhikari, Zhong-Wei Wang, Gyorgy Szabadkai, Michael R Duchen, Senthil Arumugam, Reza Nosrati, John Carroll

## Abstract

Mitochondria are highly dynamic organelles, and their spatiotemporal organization is strictly regulated. While it has long been recognized that mitochondria in ovulated oocytes are concentrated in the spindle hemisphere, the mechanism remains unknown. Through live cell imaging and modelling, we have discovered three features of mitochondrial dynamics in the ooplasm: (i) actin-driven cortical mitochondrial streaming is delimited to the boundary of the polarized spindle hemisphere; (ii) distinct from bulk cytoplasmic streaming, mitochondrial streaming occurs bilaterally and perpendicular to the long axis of the MII spindle and, (iii) mitochondria movement from the cytoplasm to the polarized cortex is via MYO19-mediated chromatin-associated channel around the spindle midzone. This directionality in mitochondrial streaming patterns the ooplasm of the spindle cortex, creating mitochondria-rich and mitochondria-poor regions. These features explain the establishment of the polar gradient of mitochondria in MII oocytes and may provide new insight into the spatiotemporal organization of mitochondria in cells.

## Introduction

Of all cellular organelles, mitochondria are the most strictly regulated in their spatiotemporal organization. Precise localization of mitochondria provides local provision of ATP and metabolites necessary to drive cellular events^1^. Synaptic activity^2^, sperm flagellar movement^3^, and lamellipodia^4^ all rely on local mitochondrial activity. In mouse oocytes, mitochondria aggregate around the developing meiotic spindle^5, 6, 7, 8, 9, 10^, presumably supporting spindle assembly and function.

This specific positioning of mitochondria is achieved via the microtubule and actin cytoskeleton. Microtubule-mediated trafficking is dominant in most cell types and is driven by kinesin- and dynein-based motor proteins coupled to mitochondria via TRAK1/2 and MIRO1/2 adaptors. Disrupting these mitochondria-specific adaptors leads to mitochondrial mislocalization and consequent aberrant cell function ^11, 12, 13, 14, 15, 16, 17, 18, 19, 20^. Mitochondrial trafficking on the actin cytoskeleton is via MYO19 ^21^, the only mitochondria-specific myosin-based motor protein. MYO19-mediated mitochondrial trafficking is a feature of metaphase cells when microtubules are subverted to the mitotic spindle, and ablation of MYO19 leads to disrupted cell division^22^.

More recently mitochondrial distribution in mitotic cells has been shown to be driven by cytoplasmic waves of actin polymerization, thereby ensuring a stochastic distribution of mitochondria to daughter cells at the time of cell division^23, 24^. The discovery of actin-driven cytoplasmic streaming in mature metaphase II (MII)-arrested oocytes^25^ raises the possibility that mitochondria are undergoing streaming-driven movement in the ooplasm. To date, cytoplasmic streaming in MII stage oocytes has been performed using particle tracking on bright field images so it is as yet unknown if mitochondria participate in streaming or if they remain anchored or trapped in the rich cytoplasmic actin network.

As oocytes progress through meiotic maturation, mitochondria undergo an extensive microtubule-mediated redistribution. From an initial relatively homogeneous cytoplasmic distribution of mitochondria around the large centrally placed germinal vesicle (GV)^6, 9^, to a final distribution in the mature Metaphase II (MII)-arrested oocyte where mitochondria are highly polarized to the hemisphere containing the meiotic spindle^5, 6^. This polarized distribution is not true of all oocyte organelles. It is well established that endoplasmic reticulum (ER) is distributed throughout the cytoplasm with increased clusters in the non-spindle hemisphere^26, 27^. The concentration of mitochondria in the spindle hemisphere serves to localize mitochondria close to the site of ATP-requiring events of chromosome segregation and the highly asymmetric meiotic divisions known as polar body extrusion (PBE), yet the mechanism underlying this polarized distribution is unknown.

Like mitotic somatic cells, the cytoplasm of the mature oocyte is bereft of microtubules except for in the large cortically located MII spindle. This suggests that mitochondrial distribution may be driven by actin-mediated trafficking^19, 25^, or by cytoplasmic actin waves and resultant cytoplasmic streaming^22^. To investigate the dynamic mechanisms underlying the gradient of mitochondria in mouse MII oocytes, we have employed live-cell imaging, molecular perturbations, and 3D computational modeling.

Our findings reveal that actin-mediated cytoplasmic streaming^25^ facilitates bulk mitochondrial movement, which we find is restricted to the spindle hemisphere of the oocyte. Remarkably, mitochondrial streaming is determined by spindle orientation and occurs perpendicular to the long axis of the spindle. This directionality is imposed by a chromosome-mediated ‘channel’ through which mitochondria pass from the sub-spindle cytoplasm to the cortex. These findings reveal a previously unappreciated complexity to mitochondrial organization in oocytes, with implications for understanding cytoplasmic organization in oocytes.

## Results

### Polarized distribution of mitochondria in MII oocytes is established by a distinct organization of mitochondrial streaming in the spindle hemisphere

We first investigated mitochondrial distribution in tetramethylrhodamine methyl ester (TMRM)-loaded MII oocytes using 2D confocal imaging. Oocytes in the field of view were imaged through a central cross-sectional plane and were readily oriented via the spindle position, as marked by the absence of TMRM fluorescence. Focusing on oocytes in which the spindle is in the plane of imaging, we noticed two main patterns of mitochondrial distribution that were associated with different spindle orientations (**Fig. 1**). First, when the oocyte is imaged with the spindle sectioned along its long axis (side-on view) mitochondria were largely aggregated in the cytoplasm immediately below the spindle (11 of 22 oocytes; **Fig. 1a,c**). Second, when the oocyte was imaged with the spindle sectioned across its orthogonal axis (end-on view), mitochondria were observed extending up either side of the spindle to the cortex and then along the cortex of the spindle hemisphere (11 of 22 oocytes; **Fig. 1b,d**). Analysis of this spindle orientation-based difference in mitochondrial distribution was performed using a fluorescence intensity profile along a line through the middle of the spindle. This demonstrates a significant difference in mitochondrial accumulation around the spindle in the two orientations (**Fig. 1e**).

**Figure. 1:**
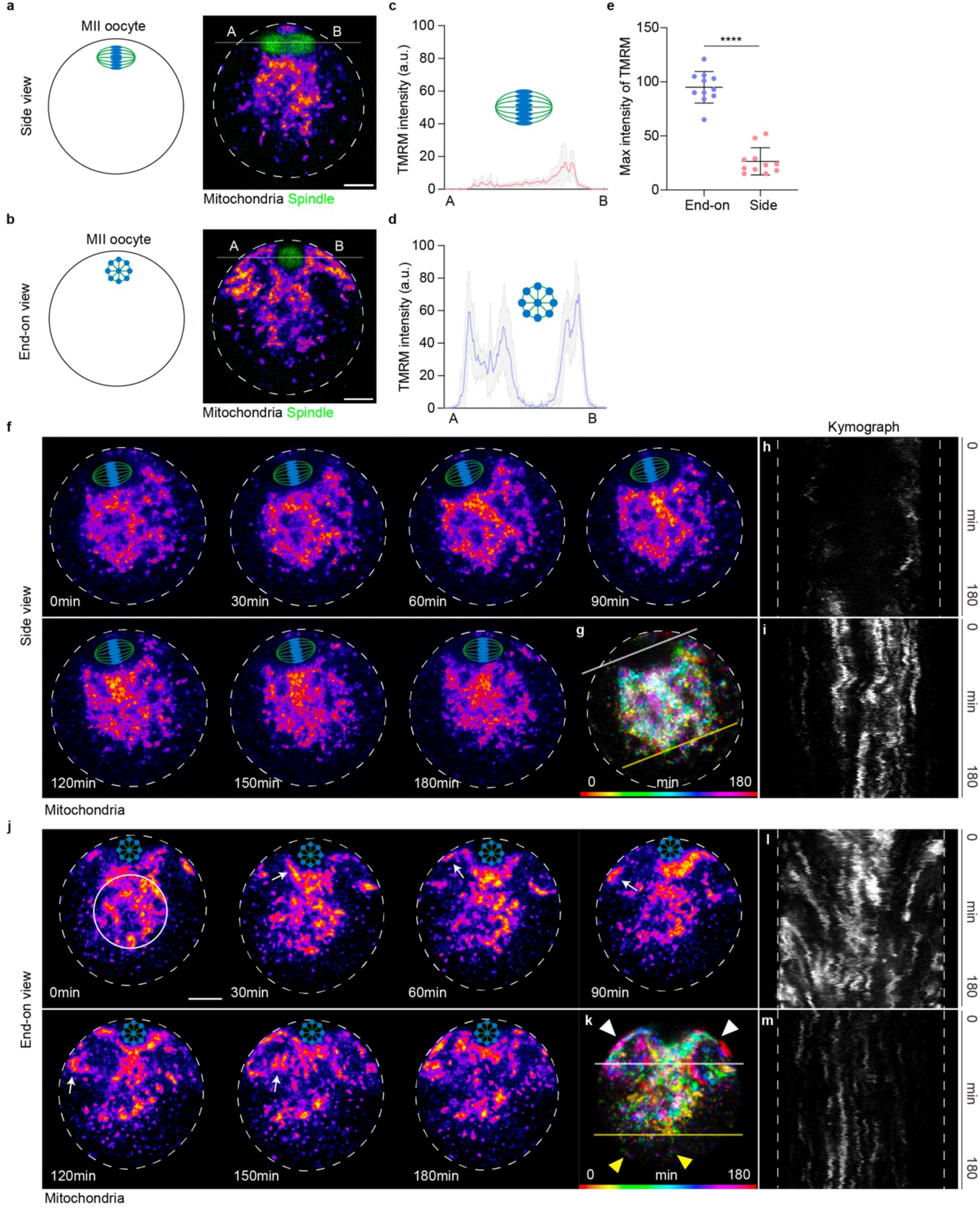
Spindle orientations define the apparent pattern of mitochondrial streaming in MII oocytes. **a,b**, Schematics illustrating the live-cell imaging for mitochondrial movement in MII oocytes and representative images showing mitochondrial distribution in the side or the end-on spindle orientations, respectively. Confocal images of a single slice covering the entire ooplasm in the middle of the spindle cross-section were observed in two orientations of the spindle axis. MII oocytes were stained with TMRM (fire) and SiR-Tubulin (green). Scale bar, 15 µm. Data represents 22 oocytes with 3 replicates. **c,d**. Measurement of TMRM intensity along a line crossing the spindle in the side or the end-on view of spindle oocytes in ‘**A’** and **’B’**. **e.** Maximal TMRM intensity measured at the line in **’A’** and **’B’**. Unpaired t-test (****p<0.0001). **f.** Mitochondrial distribution in a representative oocyte was monitored over a period of 3 hr (Δt = 30 mins) from the spindle side view. Mitochondria did not show clear bulk movement in the ooplasm. Data represents 4 oocytes with 4 replicates. **g.** Pseudo-colored time-projection image showing mitochondrial movement (Δt = 30 mins). White and yellow lines denote kymograph regions for the spindle hemisphere and non-spindle hemisphere, respectively, as shown in **h,i**. **h,i**. Kymograph analysis depicting mitochondrial movement in the spindle hemisphere (h) and non-spindle hemisphere (i) (Δt = 30 secs). Dotted line indicates the oocyte membrane. **j.** Mitochondrial distribution in a representative oocyte was observed over 3 hr (Δt = 30 mins) in the spindle end-on view. Initially, mitochondria were located below the spindle (indicated by a white circle in the first image). White arrows track the migration of mitochondrial clusters from below the spindle up to and along the cortex before redirecting toward the center of the oocyte. Dotted line indicates the oocyte membrane. Scale bar, 15 µm. Data represents 5 oocytes with 5 replicates. **k**. Pseudo-colored time-projection image showing mitochondrial movement (Δt = 30 mins). White arrowhead highlights cortical streaming mitochondria in the spindle hemisphere, contrasting with the non-spindle hemisphere (yellow arrowhead). White and yellow lines denote the kymograph regions as shown in **l,m**. **l,m.** Kymograph analysis depicting mitochondrial movement in the spindle hemisphere (l) and non-spindle hemisphere (m) (Δt = 30 secs). Dotted line indicates the oocyte membrane.

To gain further insight into this spindle orientation-associated difference in mitochondrial organization, we turned to time-lapse confocal imaging of mitochondria loaded with TMRM (**Fig 1f-m**) or mito-Dendra2 (**Supplementary Fig. 1a**). Movies of mitochondrial dynamics revealed that mitochondria imaged in the side-on view showed no directional movement (**Fig. 1f-i**, **Supplementary Video 1**), while in the end-on orientation, mitochondria were highly dynamic moving from the large aggregation of mitochondria below the spindle up either side of the spindle to the cortex (**Fig. 1j-m, Supplementary Video 1**). This pattern of mitochondrial movement was also evident in oocytes expressing mito-Dendra2 (**Supplementary Fig. 1a**). Further, as an alternative approach, mitochondria-targeted photoactivatable (PA)-GFP was expressed in oocytes and photo-converted in a small cluster of mitochondria below the spindle (**Supplementary Fig. 1b,c**, end-on view). This clearly shows photo-converted mitochondria moving up either side of the spindle to the cortex. The time course of mitochondrial movement is analyzed as a decrease in fluorescence seen in the regions of interest (ROIs) where mtPA-GFP was initially activated. Reverting to a side-on orientation and moving the plane of focus by 5-6 µm either side of the mid-plane of the spindle confirmed that mitochondria were tracking up the sides of the spindle midzone to the oocyte cortex (**Supplementary Fig. 1b,c**, side view).

Once mitochondria traverse the spindle and reach the cortex, they appear to accelerate along the cortex away from the spindle (**Fig. 1j**, white arrows). Approximately one-third of the way down the oocyte cortex, mitochondria then began tracking inward toward the cytoplasm below the spindle (**Fig. 1j**, t=120-180 min, white arrow; **Supplementary Video 1**). The dramatic difference in mitochondrial movement in the spindle hemisphere relative to the non-spindle hemisphere is further demonstrated by Time-Projection Analysis (**Fig. 1k**, white and yellow arrowheads) and kymographs of mitochondrial movement in the spindle and non-spindle hemispheres (**Fig. 1l,m**).

We then quantified the properties of mitochondrial streaming in both side-on and end-on views. As shown in **Fig. 2a**, a streamline analysis in the side-on view exhibited a largely random movement pattern. Vector component analysis along the X and Y axes confirmed this lack of directional movement of mitochondria (**Fig. 2b,c**), and the mean velocity maps further demonstrated limited mitochondrial movement: below the spindle, 0.12±0.07 µm/min (Region A in **Fig. 2d,e**); side of the spindle poles, 0.13±0.09 µm/min (Region B in **Fig. 2d,e**).

**Figure. 2:**
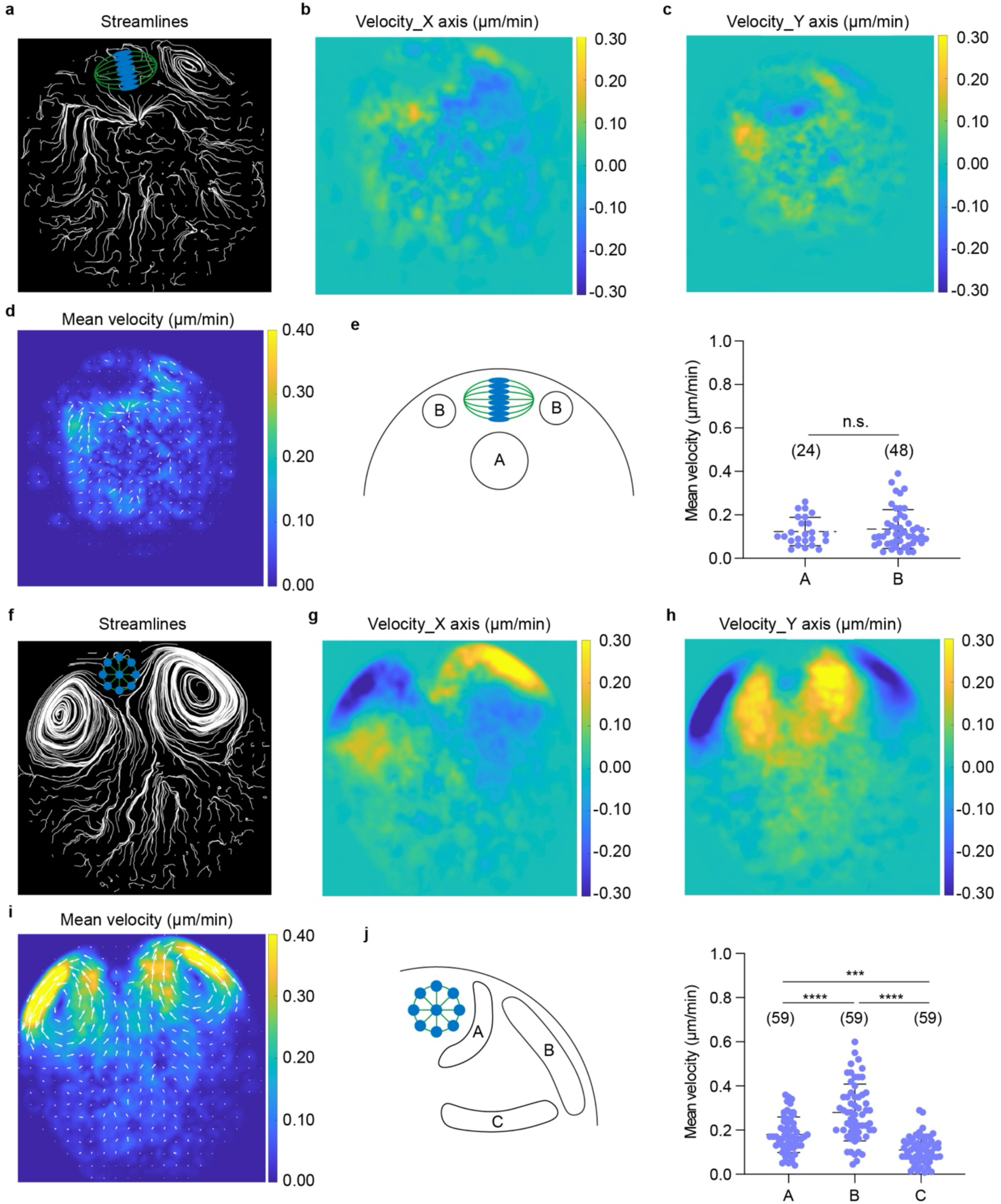
Quantification of mitochondrial streaming in the spindle end-on and side orientations. **a-d**. (**a**) Streamlines visualizing mitochondrial flow patterns in the side view of the oocyte. Decomposed velocity fields along the (**b**) X axis and (**c**) Y axis in the side view of the oocyte, and the corresponding (**d**) mean velocity field overlaid with white arrows illustrating directional variation of the velocity vectors across different regions of the oocyte during mitochondrial streaming. The color scale indicates velocity magnitude (µm/min). **e**. Quantification of velocity in different regions of the spindle hemisphere. Region A indicates below spindle and Region B indicates side of the spindle poles. Unpaired t-test (n.s.: non-significant). Analysis was undertaken on 4 individual oocytes in which mitochondrial streaming was observed in the side view. **f-i**. (**f**)Streamlines visualizing mitochondrial flow patterns in the end-on view of the oocyte. Decomposed velocity fields along the (**g**) X axis and (**h**) Y axis in the end-on view of the oocyte, and the corresponding (**i**) mean velocity field overlaid with flow arrows illustrating directional variation of the velocity vectors across different regions of the oocyte during mitochondrial streaming. The color scale indicates velocity magnitude (µm/min). **j**. Quantification of velocity in different regions of the spindle hemisphere. Region A indicates where mitochondria move from below the spindle to the cortex, Region B indicates the cortex where mitochondria slide along downward, Region C indicates where mitochondria move inwardly to cytoplasm. One-way ANOVA (***p<0.001 and ****p<0.0001). Analysis was undertaken on 5 individual oocytes in which mitochondrial streaming was observed in the end-on view.

By comparison, in the end-on spindle orientation, streamline analysis reflected the experimental observations, showing dramatic swirling patterns on either side of the spindle that were constrained to the polarized hemisphere of the oocyte (**Fig. 2f**). In contrast, mitochondrial streamlines in the non-spindle hemisphere were short and more randomly oriented. Vector component analysis along either the X or Y axis (**Fig. 2g,h**) clearly showed directionality in the pattern of mitochondrial movement in the spindle hemisphere, but not in the non-spindle hemisphere. We next determined the mean velocity in different regions of the streaming pattern: mitochondria moving upward next to the spindle traveled at 0.18±0.08 µm/min (Region A in **Fig. 2j**); accelerated to 0.28±0.13 µm/min as they progressed along the cortex (Region B in **Fig. 2j**); before slowing to 0.11±0.06 µm/min (Region C in **Fig. 2j**) when leaving the cortex and moving toward the sub-spindle cytoplasm (**Fig. 2i,j**). This sharp change in velocity and direction of mitochondrial streaming occurred at the boundary of the polarized spindle cortex, as indicated by Concanavalin A labelling (**Supplementary Fig. 2**). This spindle hemisphere restriction of mitochondrial streaming provides an underlying mechanism that explains the well-established polarized accumulation of mitochondria in MII oocytes.

Our previous studies on maturing oocytes reported that spindle-associated mitochondria have a higher MMP than those in the peripheral cytoplasm, so we asked if there is any activity-dependent gradient of mitochondria across the oocyte. To test this, we used a ratiometric technique in which TMRM fluorescence is ratioed against an MMP-independent indicator, mito-Dendra2^28^. This analysis showed that MMP was significantly higher in mitochondria located in the cytoplasm immediately below the MII spindle compared to those in the non-spindle hemisphere (**Supplementary Fig. 3a**). Further, this difference was not seen when the ratiometric analysis was performed using two MMP-independent indicators, mito-mScarlet and mito-Dendra2 (**Supplementary Fig. 3b**). Thus, it appears not only are more mitochondria localized to the spindle hemisphere, but they are also more active.

### Actin polymerization drives mitochondrial streaming and accumulation in the cortical hemisphere

The phenomenon of cytoplasmic streaming, first reported by Yi et al. (2011) is driven by actin polymerization in the polarized spindle cortex. To begin investigating the relationship between mitochondrial streaming and cytoplasmic streaming, we have disrupted cytoskeletal dynamics using nocodazole, to inhibit microtubule polymerization, and cytochalasin D (CCD), to inhibit actin polymerization (**Fig. 3a-c**). Nocodazole treatment had no noticeable effect on mitochondrial streaming (**Fig. 3b**), while inhibition of actin polymerization completely blocked mitochondrial movement (**Fig. 3c**) as evidenced by the large aggregation of mitochondria in the sub-spindle cytoplasm and analysis of the timeseries projection in control and CCD-treated oocytes (**Fig. 3a-e**). Inhibition of mitochondrial streaming by CCD treatment led to a loss of mitochondrial accumulation in the spindle hemisphere, while control and nocodazole-treated oocytes retained the expected polarized mitochondrial distribution (**Fig. 3f,g**). Thus, actin polymerization and subsequent mitochondrial streaming are necessary to establish and maintain the polarized distribution of mitochondria in MII oocytes.

**Figure. 3:**
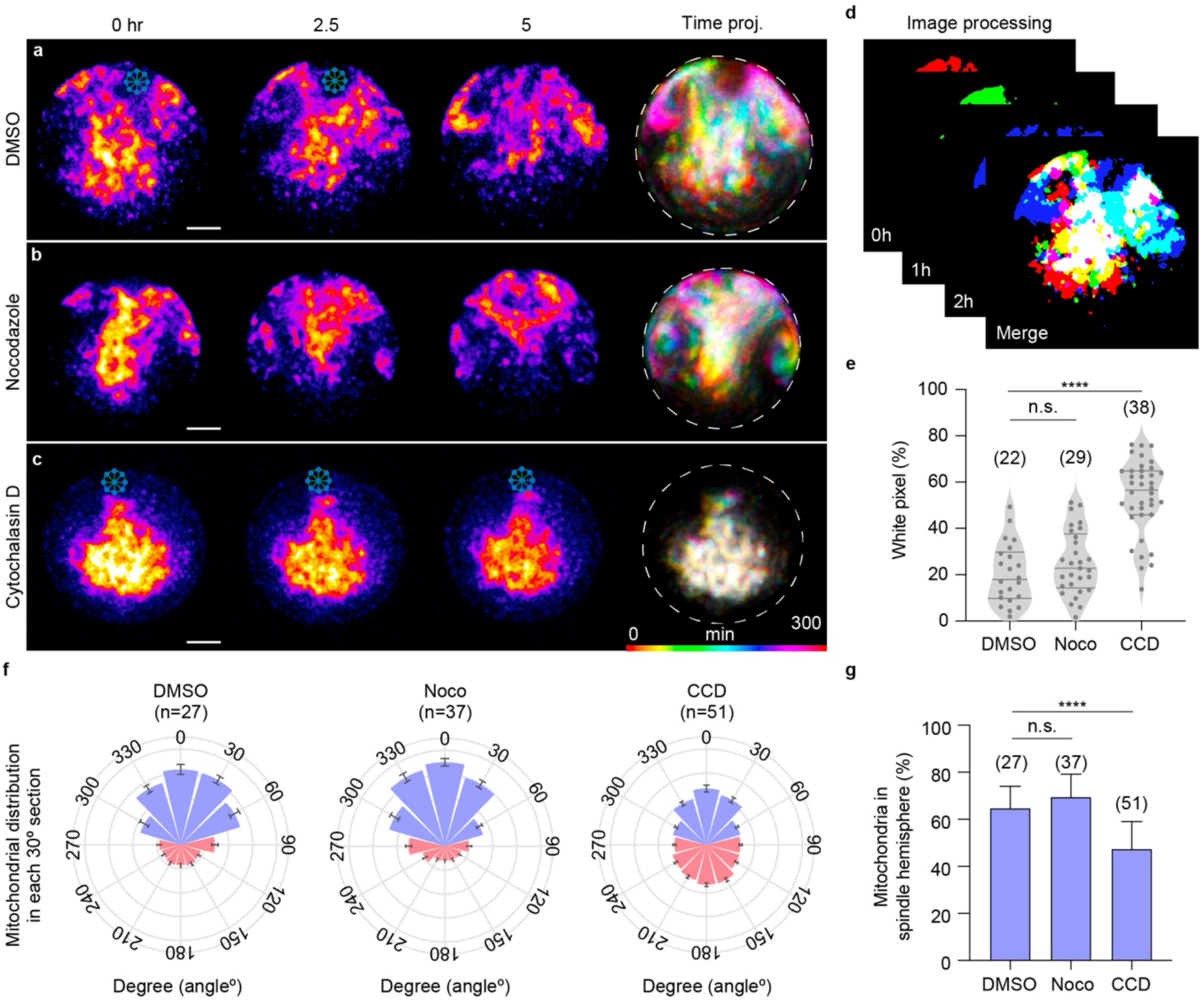
Actin cytoskeleton drives bulk mitochondrial streaming. **a-c,** Representative images showing mitochondrial streaming in oocytes treated with DMSO (**a**), nocodazole (**b**), or CCD (**c**). Time-projection images with spectral pseudo-colors are presented at the end of each row (Δt = 3 mins). At the start of imaging, spindles were positioned at the 12.00 o’clock position. In nocodazole-treated oocytes, chromosomes disperse towards the cortex as microtubules depolymerize. Mitochondria were labeled with TMRM. Dashed line indicates the oocyte membrane. Scale bar, 15 µm. Data represents 22-38 oocytes from 3 replicates. **d,** Schematic representation of pixel quantification analysis. Mitochondrial clusters were segmented at 0, 1, and 2 hr and pseudo-colored red, green, and blue, respectively. The time-merged image displayed seven colors according to pixel alignment: red, green, blue, yellow, purple, cyan, and white. **e,** Quantification of percentage of white pixels in oocytes treated with DMSO, nocodazole, or CCD. Note that a higher percentage of white pixels indicates less mobile mitochondria over time. Unpaired t-test (****p<0.0001 and n.s.: non-significant). **f,** Quantification of mitochondrial populations in the spindle hemisphere versus the non-spindle hemisphere in oocytes treated with DMSO, nocodazole, or CCD. Each oocyte was divided into 12 30-degree sections. Only oocytes with the spindle positioned at 12 o’clock position were analyzed. Blue represents mitochondria in the spindle hemisphere (150 degrees), while red indicates those in the non-spindle hemisphere (210 degrees). **g,** Percentage of mitochondria in the spindle hemisphere relative to total mitochondrial population measured in **f**. Data are presented as mean ± SD. Unpaired t-test (****p<0.0001 and n.s.: non-significant).

### Chromatin coordinates mitochondrial and cytoplasmic streaming

During treatment with nocodazole, chromosomes scatter around the cortex in an actin-dependent manner and cause local cortical differentiation as marked by increased actin polymerization^29^ (**Supplementary Fig. 4**). Imaging mitochondria in the region of isolated chromatin revealed that local mitochondrial streaming events occur around each of the chromatin clumps (**Fig. 4a, Supplementary Video 2**). Particle Image Velocimetry (PIV) analysis confirmed that, as for mitochondria, cytoplasmic particles tracked toward each dispersed chromatin clump in all nocodazole-treated oocytes examined (**Fig. 4b,c**). The velocity of particle movement in these local regions was reduced in nocodazole-treated oocytes compared to controls, presumably due to the lower concentration of chromatin and relative amount of actin polymerization at these sites.

**Figure. 4:**
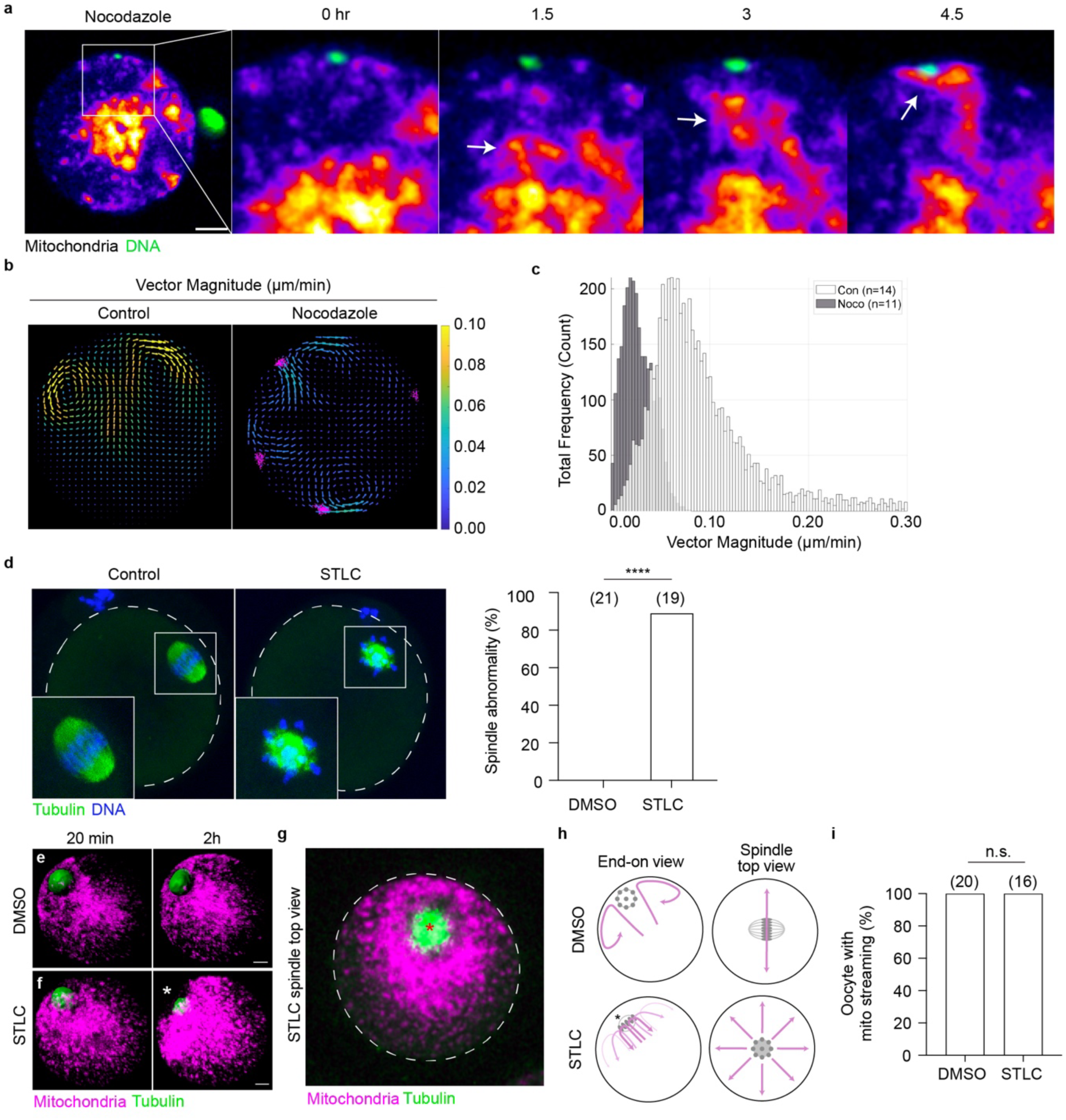
Chromosome alignment determines the directionality of mitochondrial streaming. **a,** Representative images showing mitochondrial streaming in nocodazole-treated oocytes with mitochondria labeled by TMRM and chromosomes stained with SYBR Green. Note that mitochondrial clusters migrate toward dispersed chromosomes (indicated by arrow). Data represents 37 oocytes from 3 replicates. Scale bar, 15 µm. **b**, PIV analysis of control oocytes in the end-on spindle view and nocodazole-treated oocytes. Control oocytes (n=7) show two strong swirling streaming patterns at either side of the spindle. In nocodazole-treated oocytes (n=4), multiple relatively smaller flows are observed around dispersed chromosomes (magenta) near the cortex. **c**, Quantification of vector magnitude in the swirling regions of control oocytes (each side of the spindle, indicated by warm-colored arrows) and nocodazole-treated oocytes (each dispersed chromosome). **d,** Representative images of spindles and quantification of abnormal spindle percentage in DMSO- or STLC-treated oocytes. Chi-square test (****p<0.0001). Images represent 19-21 oocytes from 3 replicates. **e,f,** Representative 3D projections showing mitochondrial distribution in oocytes treated with DMSO (**e**) or STLC (**f**) after 20 min and 2 hr. Mitochondria are labeled with TMRM, and spindles with SiR-Tubulin. Time point 0 corresponds to the first imaging frame after STLC addition. Asterisk indicates a mitochondria-free region at the cortex. Δz = 1.5 µm. Scale bar, 10 µm. Images represent 16-20 oocytes from 3 replicates. **g,** Representative of 3D projected mitochondrial distribution in STLC-treated oocytes observed from the spindle top view. Asterisk indicates the mitochondria-free region at the cortex. Dashed line indicates the oocyte membrane. Δz = 1.5 µm. Images represent 16 oocytes from 3 replicates. **h,** Schematic illustrations of mitochondrial movement in DMSO- or STLC-treated oocytes from different views. Magenta arrows indicate mitochondrial movement and directionality. Asterisk indicates the mitochondria-free region at the cortex. **i,** Percentage of oocytes with mitochondrial streaming patterns following STLC treatment. Chi-square test (n.s.: non-significant)

Next, we sought to perturb the organization of chromatin in the oocyte to further test its role in the spatiotemporal patterning of mitochondria in oocytes. First, we created a chromatin ‘rosette’ in the cortex by using the Eg5 inhibitor, S-Trityl-L-cysteine (STLC), to generate a monopolar spindle (**Fig. 4d**). In the presence of this chromosome configuration, mitochondrial streaming was no longer bilateral but occurred in a radial ‘fountain’-like pattern around the chromosomes (**Fig. 4e-i, Supplementary Video 3,4**). Second, we used micromanipulation approaches to remove the MII spindle entirely or to transplant it to the non-spindle cortex or to a different MII oocyte. In spindle-enucleated oocytes, mitochondrial streaming was abolished (**Supplementary Fig. 5, Supplementary Video 5**), while in reconstructed oocytes mitochondrial streaming tracked the new spindle location (**Supplementary Fig. 6,7, Supplementary Video 6,7**).

Chromatin-mediated actin polymerization^30, 31^ and cytoplasmic streaming^25^ are known to be driven by Ran-GTP. To confirm if Ran-GTP-dependent actin polymerization is also driving mitochondrial streaming, we injected mRNA encoding dominant negative Ran T24N to inhibit cortical actin formation (**Supplementary Fig. 8a-c**). Strikingly, Ran T24N-injected oocytes showed a mitochondrial distribution phenotype similar to that seen after actin inhibition by CCD: mitochondria remained localized in the central cytoplasm (**Supplementary Fig. 8d**); there was no evidence of mitochondrial streaming (**Supplementary Fig. 8e**); and mitochondria were distributed evenly across both hemispheres (**Supplementary Fig. 8f,g**). Control oocytes injected with WT Ran mRNA demonstrated normal mitochondrial streaming and a spindle hemisphere-oriented mitochondrial distribution (**Supplementary Fig. 8d-g**). Together, these experiments confirm that, like cytoplasmic streaming, the pattern of mitochondrial movement is coordinated by the location of the chromatin, and that Ran-GTP emanating from the chromatin drives this movement through local cortical actin polymerization.

### Chromatin-mediated channeling of mitochondria at the spindle midzone determines the pattern of mitochondrial streaming

The role of chromatin in coordinating mitochondrial streaming led us to ask if chromatin on the MII spindle is involved in establishing the mitochondrial distribution seen in different spindle-orientations. To more fully appreciate mitochondrial dynamics around the spindle, a 3D analysis of mitochondrial distribution was performed. As could be predicted from the 2D images, the 3D end-on view shows mitochondria in abundance around the sides of the spindle and at the cortex (**Fig. 5a**, end-on view). Remarkably, rotation of the same oocyte to visualize the side-on view in 3D revealed a band of mitochondria tightly aligned with the position of the chromosomes in the midzone of the MII spindle (**Fig. 5a**, side view, white arrow). This organization was further demonstrated in individual sections and the projection image of 7 sections through the spindle (**Fig. 5b**) and was seen in all 13 oocytes examined (**Supplementary Video 8**). Analysis of fluorescence intensity of microtubules, DNA, and mitochondria along the long axis of the spindle (**Fig. 5c**) demonstrates that mitochondria are tightly aligned with the chromosomes on the metaphase plate.

**Figure. 5:**
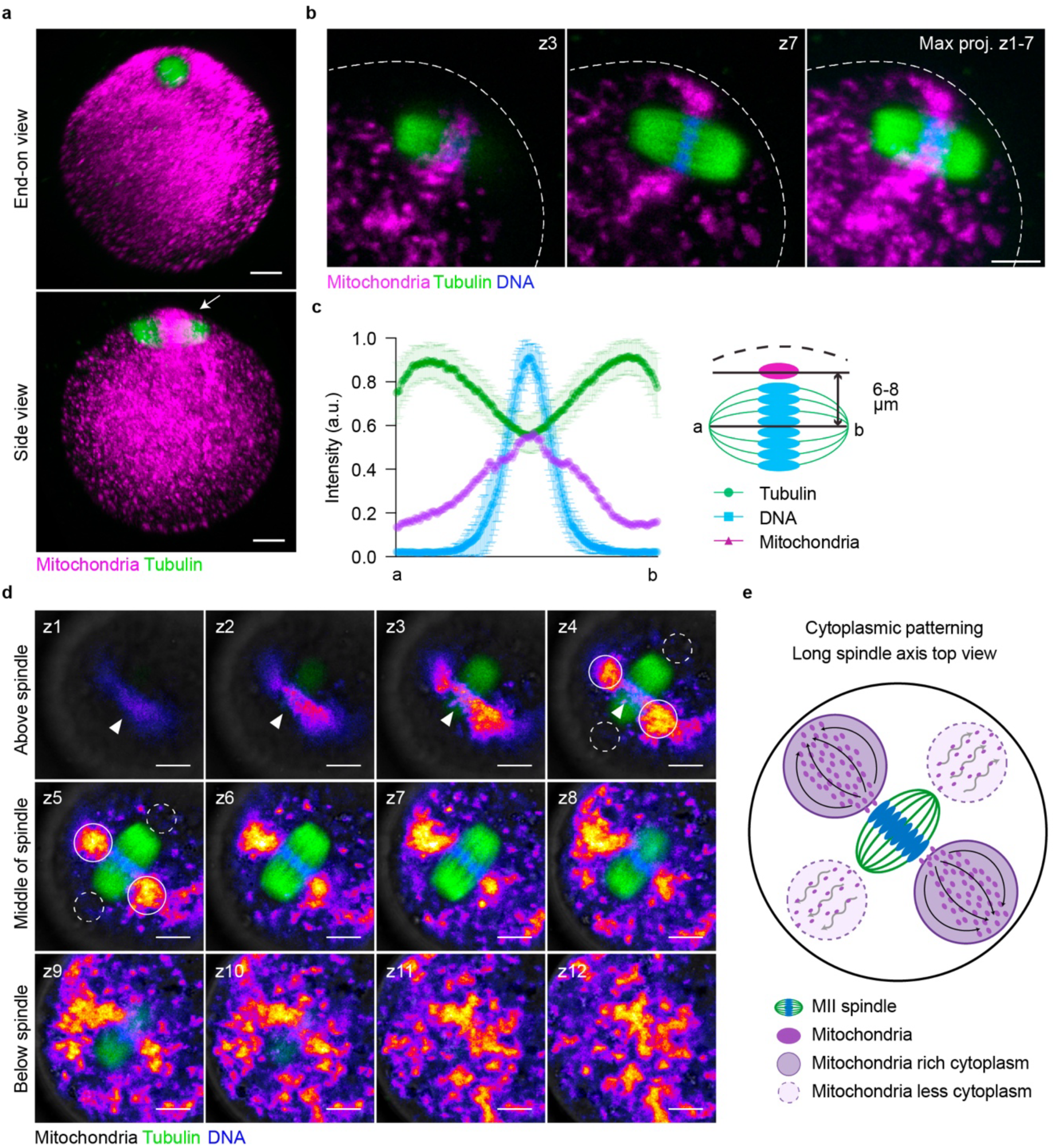
Chromatin-mediated channeling of mitochondria at the spindle midzone determines the pattern of mitochondrial streaming. **a,** Representative 3D projections of oocyte mitochondria (TMRM) and tubulin (SiR-Tubulin). Note that in the end-on view, mitochondria fill the cortex perpendicular to the spindle, while in the side-on view the cortical area shows limited mitochondrial density. Mitochondria are seen wrapping around the spindle midzone (white arrow). Δz = 1.5 µm. Scale bar, 10 µm. Note that this distinct mitochondrial distribution at the different spindle orientations was observed in all oocytes exhibiting mitochondrial streaming in the spindle hemisphere. **b,** Representative single-plane and projection images of oocyte mitochondria (TMRM), tubulin (SiR-Tubulin), and DNA (Abberior DNA dye). Note the close alignment of mitochondria and chromosomes. Dashed lines indicate the oocyte membrane. Δz = 2 µm. Scale bar, 10 µm. Images represent 29 oocytes from 2 replicates. **c**, Schematic and quantification of TMRM, Abberior DNA dye, and SiR-Tubulin intensities. Tubulin and DNA intensities were measured at the spindle midzone (black line in the spindle schematic), while TMRM intensity was measured in the region between the cortex and the spindle (black line passing through the magenta oval). The peak of TMRM intensity aligns with the DNA intensity peak. 22 oocytes analyzed. **d,** A series of z-slices taken along the long axis of the spindle, starting from above the spindle (z1-3), through the spindle (z4-8), to the ooplasm below the spindle (z9-12). Fire LUT, mitochondria (mito-Dendra2); green, spindle (SiR-Tubulin); blue, DNA (Abberior DNA dye). White arrowheads indicate mitochondrial distribution immediately above the spindle. White circles on z5 indicate mitochondrial-rich regions neighboring the midzone, while dashed circles denote mitochondrial-sparse areas in the ooplasm parallel to the spindle long axis. Δz = 1.8 µm. Data represents 29 oocytes from 2 replicates. Scale bar, 10 µm. **e,** Schematic depiction of the cytoplasmic patterns established by chromatin-induced directionality of mitochondrial streaming. Arrow thickness and color reflect mitochondrial flow intensity, with thicker and darker arrows indicating stronger movement.

The data thus far indicate that the spindle orientation-determined mitochondrial distribution results from a preferred chromosome-mediated ‘channel’ through which mitochondria move from cytoplasm below the spindle to above the cortex. This organization confers a directionality to mitochondrial streaming in the spindle hemisphere such that the flow occurs bilaterally and perpendicular to the long axis of the spindle. An inevitable outcome of this directionality is a previously unappreciated spatial patterning of the ooplasm of the spindle hemisphere, whereby the ooplasm perpendicular to the spindle is abundantly populated by mitochondria and presumably other organelles involved in streaming, while ooplasm extending parallel to the spindle poles is relatively quiescent and with fewer mitochondria (**Fig 5d,e; Supplementary Video. 9**).

### Modeling fluid dynamics in the ooplasm

To determine if cytoplasmic fluid dynamics alone is sufficient to generate the orientation-dependent streaming patterns and the precise mitochondrial channeling at the spindle midzone, or if additional active mechanisms are required, we developed a 3D Stokes-flow model. We investigated the relationship between the observed mitochondrial streaming pattern and the forces generated either at the center of oocytes or at the cortex and explored three predicted scenarios.

First, we applied a localized volumetric force at the cytoplasm below the spindle toward the cortex, mimicking the initiation of bulk mitochondrial movement (Scenario A, **Fig. 6a**). While this produced a strong upward flow and weak circulation on both sides of the spindle, the simulated end-on and side-on views remained nearly identical (**Fig. 6d**), failing to replicate the strong orientation bias observed in our experiments (**Fig. 2**). Second, we modeled a uniform cortical cap driving tangential slip around the spindle (Scenario B, **Fig. 6b**). Although this generated the prominent vortex pairs seen in the end-on view, it also produced strong vortices in the side view (**Fig. 6e**), which does not match our experimental data. Finally, we introduced anisotropic cortical slip, where the force is strongest perpendicular to the spindle axis and reduced along its length (Scenario C, **Fig. 6c**). This successfully reproduced both the distinct vortices in the end-on view and the marked attenuation of in-plane flow in the side-on orientation (**Fig. 6f**). Actin polymerization, if not bilaterally restricted, has been shown to induce a flow field as in Scenario B ^25, 32^. However, bilaterally restricting mitochondria access to the cortical flow could yield similar flow dynamics as described for the anisotropic slip parallel to the spindle in Scenario C. Hence, these results indicate a non-uniform velocity field with a dominant mitochondrial streaming channel perpendicular to the spindle major axis, which is absent in the side-view plane.

**Figure. 6:**
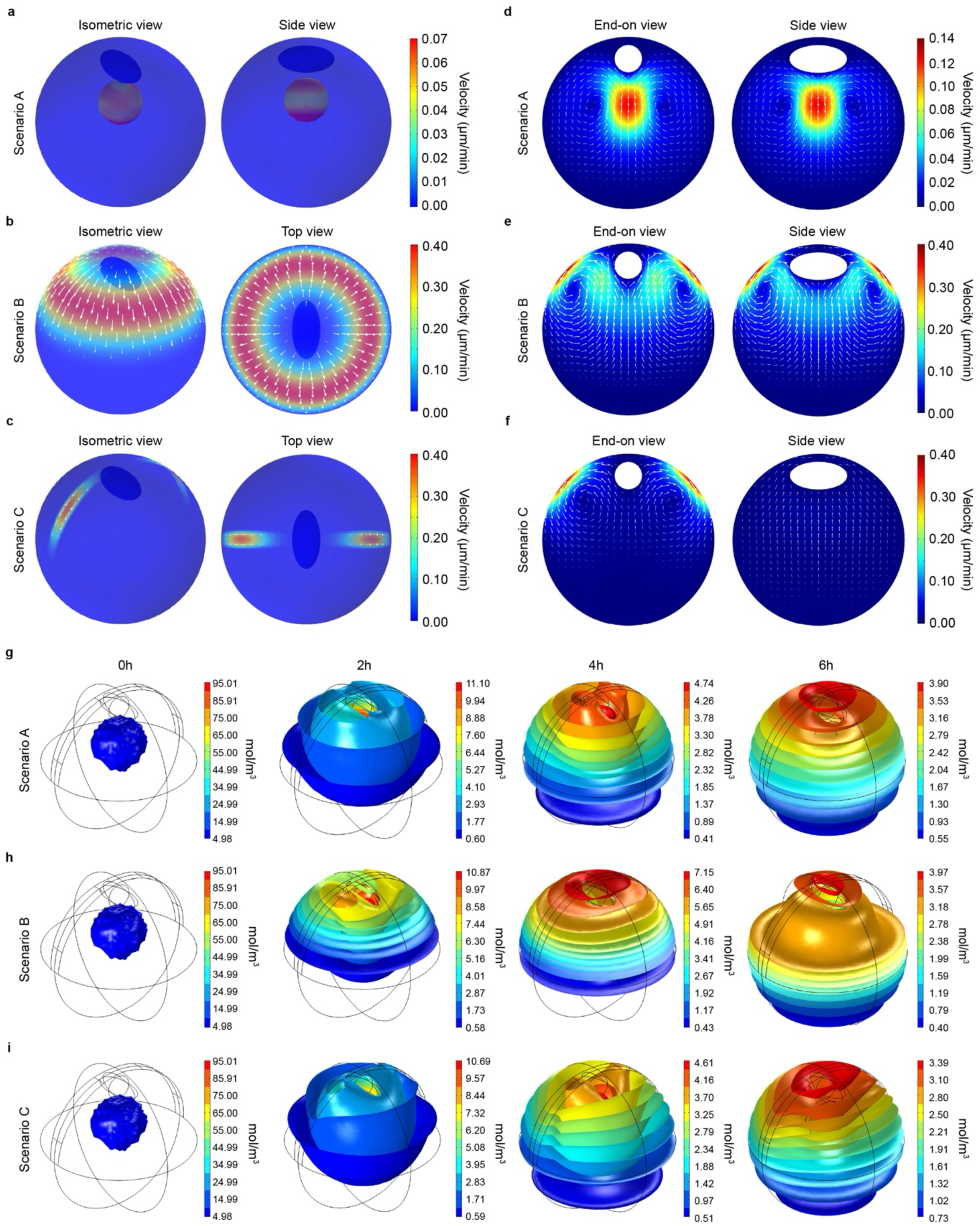
Numerical simulation of cytoplasmic fluid dynamics scenarios and tracer redistribution. **a-c**. Schematic representations of the three simulation scenarios in a 3D model of a mouse oocyte (75 µm diameter) with the spindle (dark blue ellipsoid). **a**. Scenario A: localized volumetric force applied below the spindle directed toward the cortex. **b**. Scenario B: uniform tangential slip velocity imposed on the cortical cap overlying the spindle. **c**. Scenario C: anisotropic tangential slip velocity on the cortical cap with forcing strongest perpendicular to the spindle major axis. **d-f**. Computed steady-state velocity fields for each scenario shown in end-on and side views. Color scales represent velocity (µm/min), white arrows indicate flow direction and magnitude. **d,** Scenario A shows symmetry between views but lacks the experimental vortex structure. **e,** Scenario B generates strong vortices in both planes. **f**. Scenario C reproduces the experimental observation of a strong end-on vortex pair with significantly attenuated flow in the side view. **g-i**. Time-course of passive tracer redistribution (simulating mitochondrial transport) for Scenario A, B, and C. Tracer is initially localized in a sub-spindle volume. Color scales indicate tracer concentration (mol/m^3^). **g,** Scenario A and **h,** Scenario B results in broad, hemispherical spreading of the tracer. **i,** Scenario C shows more focused accumulation near the spindle due to anisotropic forcing. However, all three hydrodynamic models fail to achieve the sharp midzone confinement observed experimentally.

We next asked whether flow geometry alone could explain the spindle midzone mitochondrial channeling. For each case, we solved an advection–diffusion problem for a passive tracer, initially localized beneath the spindle, to approximate mitochondrial redistribution by computed fluid dynamics (see Materials and methods). In both Scenario A and B (**Fig. 6g,h**), tracers were delivered into the spindle hemisphere contacting the spindle and cortex, however, their distributions remained broad, spreading over an extended band rather than forming a narrow midzone corridor. In Scenario C (**Fig. 6i**), while anisotropic cortical forcing produced a more focused accumulation in the spindle hemisphere, tracer enrichment still extended well beyond the midzone regions. Thus, passive hydrodynamics in all three models reproduced hemisphere-scale delivery but did not, by itself, generate the sharp midzone confinement seen in experimental images. These computational results demonstrate that while anisotropic cortical actin flow is sufficient to drive the global topology of mitochondrial streaming, hydrodynamics alone is insufficient to explain the localized spatial precision of the midzone channel. This suggests that the distinct spindle midzone restriction of mitochondria requires an additional level of active regulation beyond bulk cytoplasmic flow.

### The spindle midzone ‘channeling’ of mitochondria is MYO19 dependent

Given that mitochondrial movement in mitotic cells is regulated by actin rather than microtubules^22, 23, 33^, we hypothesized that this active mechanism may be due to actin-mitochondria interactions. To investigate this, we imaged mitochondria in the spindle region of oocytes expressing the actin sensor, GFP-UtrCH^34, 35^. This confirmed actin was widely distributed in and around the spindle^36^ and revealed mitochondria-actin interactions below the spindle were only apparent at the spindle midzone (**Fig. 7a,b** yellow arrows, **Supplementary Video 10**) where mitochondria are concentrated relative to the rest of the spindle (**Supplementary Fig. 9**). The presence of actin-mitochondria interactions specifically in the midzone and the absence of mitochondria along the length of the spindle points to an active actin-mediated mechanism drawing mitochondria from the cytoplasm to the spindle midzone (**Supplementary Fig. 9**).

**Figure. 7:**
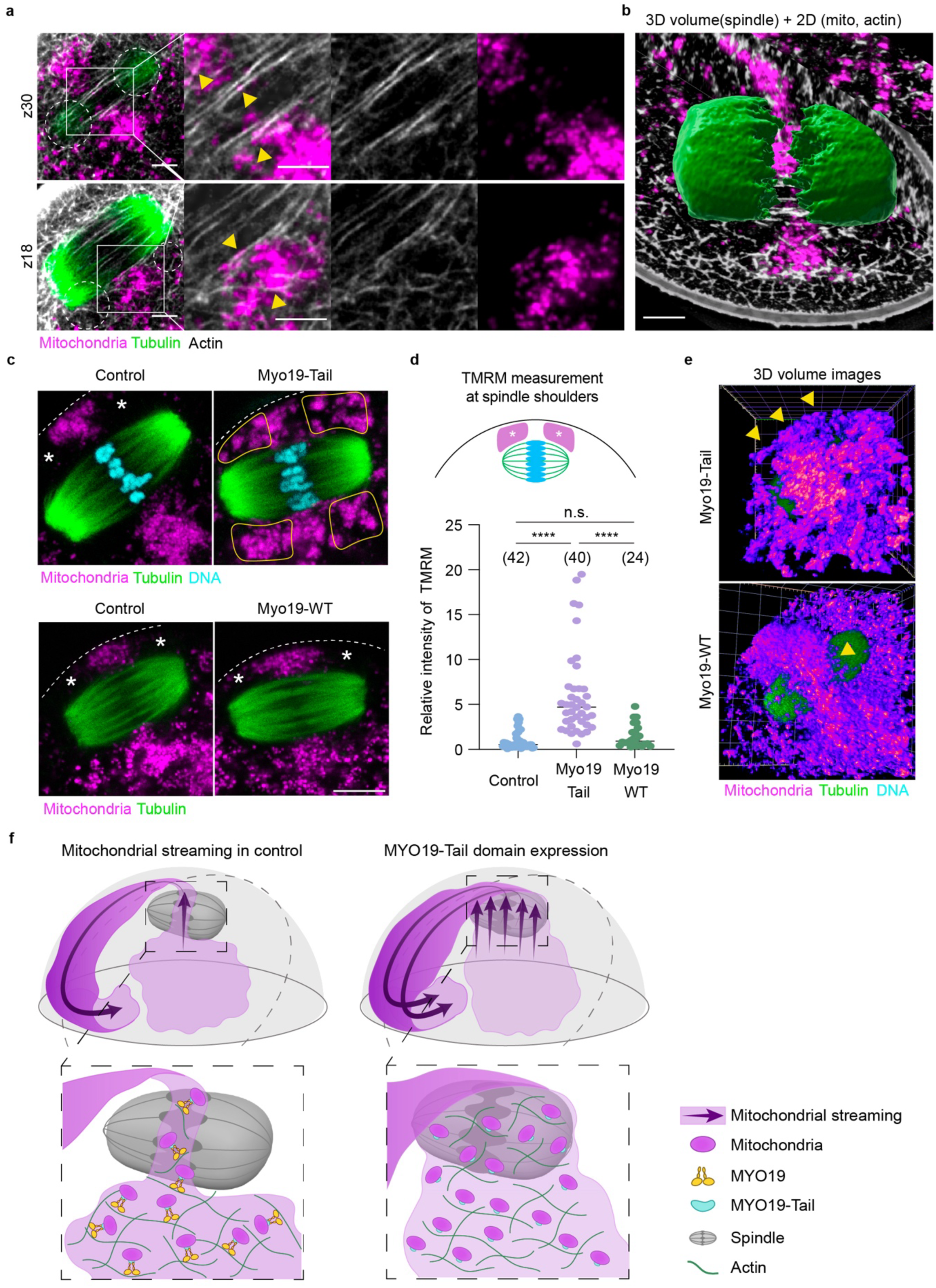
MYO19 mediates mitochondrial channeling at the spindle midzone. **a,** Representative 2D image of mitochondria (TMRM), tubulin (SiR-Tubulin), and actin (GFP-UtrCH). Yellow arrowheads indicate mitochondria aligning with actin filaments at the midzone. Dashed circle marks the area outside the midzone, where mitochondria are absent. Δz = 0.5 µm. Scale bar, 5 µm. Data represents 10 oocytes from 2 replicates **b,** Representative 3D projection of mitochondria (TMRM), tubulin (SiR-Tubulin), and actin (GFP-UtrCH) (same oocyte as in **a**). Mitochondria can be seen tightly traversing the spindle midzone. Δz = 0.5 µm. Scale bar, 5 µm. Data represents 10 oocytes from 2 replicates. **c,** Representative images showing labeled mitochondria (TMRM), tubulin (SiR-Tubulin), and DNA (SPY-DNA) after *Myo19*-Tail or *Myo19*-WT expression. Note the spindle midzone channel of mitochondria is no longer present in *Myo19*-Tail injected oocytes, resulting in mitochondria enveloping the full length of the spindle. Asterisks indicate mitochondrial-free domains between the cortex and the spindle, while yellow-outlined regions indicate mitochondria distributed between the cortex and the spindle. Dashed line indicates the oocyte membrane. Scale bar, 10 µm. Images are representative of 24-42 oocytes from 3 replicates. **d,** Schematic showing the locations of ROIs used to measure mitochondrial fluorescence intensity (top), and corresponding quantification (bottom). Note that *Myo19*-Tail-injected oocytes show increased mitochondrial presence in the spindle shoulder regions. Unpaired t-test (****p<0.0001). **e,** Representative 3D volumetric images showing mitochondrial distribution (TMRM) following *Myo19*-Tail overexpression. Yellow arrowheads indicate mitochondria located between the oocyte cortex and the spindle. In *Myo19*-WT expressing oocytes, the mitochondrial channel is prominent, whereas in *Myo19*-Tail expressing oocytes, mitochondria surround the entire spindle. Data are representative of 10-12 oocytes per group. Δz = 1 µm. **f,** Schematic model of MYO19-mediated mitochondrial channeling. In controls, MYO19 tethers mitochondria to the sub-spindle actin network, restricting their participation in bulk cytoplasmic streaming. At the spindle midzone, MYO19 facilitates active transport of mitochondria through a narrow chromatin-adjacent channel to the cortex. MYO19-Tail domain expression uncouples mitochondria from actin, allowing passive movement around the entire spindle length in bulk cytoplasmic streaming.

Actin-mitochondria interactions in somatic cells have been shown to be mediated via MYO19 ^21, 22^. To investigate a possible role of MYO19 in mitochondrial organization in MII oocytes we first confirmed it was present and colocalized to mitochondria. MYO19 immunofluorescence images showed a distinct punctate distribution that localizes with mito-Dendra2-labelled mitochondria, the bulk of which are localized to the sub-spindle cytoplasm (**Supplementary Fig. 10**). Next, we asked if MYO19 plays a role in mediating the restricted mitochondrial distribution at the spindle midzone. We compared mitochondrial distribution in oocytes injected with a dominant negative MYO19-Tail domain^21^, which contains the mitochondrial binding domain but not functional motor domains, with oocytes injected with full-length MYO19, and uninjected controls. First, we confirmed that the bulk mitochondrial streaming pattern was not affected by expressing *Myo19*-Tail domain or full-length *Myo19* (**Supplementary Fig. 11**). With no observable effect of manipulating MYO19 on mitochondrial streaming, we analyzed the distribution of mitochondria around the MII spindle. Strikingly, expression of the *Myo19*-Tail domain led to a significant increase in accumulation of mitochondria along the length of the spindle that was absent in uninjected controls or in oocytes injected with full-length *Myo19* (**Fig. 7c-e**). These findings suggest that disrupting actin-mitochondria interactions via MYO19-Tail domain leads to mitochondria moving to the cortex via the entire length of the spindle rather than the restricted spindle midzone. We therefore suggest a model whereby (i) MYO19 restrains the bulk of mitochondria on an actin network in the sub-spindle cytoplasm, thereby limiting participation in flow to the cortex, and (ii) a chromatin-and MYO19-mediated mechanism that facilitates a narrow path for mitochondria to traverse the chromatin to reach the cortical actin flow.

## DISCUSSION

The mature oocyte contains approximately 200,000 individual mitochondria, and the majority are localized in the polar spindle cortex. Here, through the use of imaging, experimental interventions, and computational modeling, we demonstrate how this polarized mitochondrial distribution is established, and reveal a chromatin-mediated mitochondrial path to the polar cortex that imposes a directionality to mitochondrial streaming in the spindle hemisphere. Further, this combination of bulk flow combined with sub-cellular limitations leads to a spindle-orientation-dependent patterning of the cytoplasm in the spindle hemisphere.

Live-cell imaging of labeled mitochondria has revealed that mitochondria in MII oocytes are highly dynamic cytoplasmic organelles that would appear to participate as a component of bulk cytoplasmic streaming first described by Yi et al. (2011). Both mitochondrial and cytoplasmic streaming are driven by and absolutely reliant on Ran-GTP-mediated cortical actin polymerization. They also have similar order of magnitude velocities, although mitochondrial streaming is approximately 40% slower than reported for cytoplasmic particles (0.28±0.13 µm/min and 0.47±0.05 µm/min, respectively). This difference is most likely explained by PIV analysis being performed in UtrCH-expressing oocytes, which we find dramatically increases cortical actin polymerization and therefore may drive an increase particle velocity. However, it is also possible that mitochondria experience increased friction due to their relatively large size compared to the particles identified in PIV/STICS analysis^37^. The dual reliance of mitochondrial and cytoplasmic streaming on Ran-GTP and actin polymerisation, as well as the similar order of magnitude velocities, strongly suggest that mitochondria in the cortex join the bulk flow arising from the polarized actin cap. However, our focus on tracking mitochondria rather than cytoplasmic particles ^25, 38, 39, 40^ has revealed a number of novel findings that provide new insights into subcellular cytoplasmic organization in oocytes.

The visualization of mitochondrial streaming and subsequent analysis of streamline vortices shows mitochondrial streaming, unlike bulk cytoplasmic streaming^25^ is (i) restricted to the spindle hemisphere and (ii) oriented perpendicular to the axis of the spindle. The abrupt cessation of cortical mitochondrial flow and subsequent inward movement at the border of the actin cap is what constrains mitochondrial distribution to the spindle hemisphere, thereby explaining this fundamental property of oocyte mitochondrial distribution^5^.

It has been suggested previously that the concentration of mitochondria to the spindle hemisphere is important for providing the energy and metabolites needed for spindle function and polar body formation^7, 10, 41, 42, 43^. Our experiments reveal that not only are mitochondria concentrated in the polar hemisphere, they also have a higher MMP and are therefore more active than those in the non-spindle hemisphere. We have previously shown in maturing oocytes where the MI spindle forms in the central cytoplasm, that spindle-associated mitochondria have a higher MMP than those in the peripheral cytoplasm^28^. The subsequent migration of the MI spindle and associated mitochondria to the cortex provide a potential explanation for how this gradient in mitochondrial MMP is first established. The functional significance of the MMP gradient remains to be determined, but local ATP production and/or supply of metabolites such as NAD+ may be important factors for subsequent events of oocyte activation and early development^7,41, 42, 44^.

The spindle-orientation dependence of mitochondrial streaming results in a bilateral flow perpendicular to the long axis of the spindle. One consequence of this asymmetry is that cytoplasmic organization differs relative to spindle orientation. Ooplasm perpendicular to the spindle is abundantly populated by mitochondria and presumably other organelles caught up in cytoplasmic vortices, while the ooplasm extending parallel to the spindle poles is relatively quiescent with limited mitochondria (see **Fig. 5d,e; Supplementary Video. 9**). The functional relevance of this cytoplasmic patterning is not clear, but it may be that centripetal forces generated by cytoplasmic vortices perpendicular to spindle orientation promote spindle stability^25^, as well as ensure spindle rotation and subsequent elongation associated with chromosome segregation occurs in the plane of polar body extrusion^38, 40^. We anticipate these observations will initiate further modeling to test these hypotheses.

The finding that mitochondria travel to the cortex via a narrow channel of chromatin-adjacent cytoplasm provides the physical restraint needed to impose directionality to mitochondrial streaming. This is supported by our modeling data which indicates neither a localized volumetric force from below the spindle, nor a uniform cortical actin cap providing cortical flow around the spindle, recapitulate the spindle-orientation-dependent streaming patterns observed experimentally. Only when the flow was constrained to a bilateral anisotropic cortical force, which effectively replicates the scenario in which access of mitochondria to the cortex is bilaterally restricted, did the model successfully reproduce the asymmetry in the mitochondrial vortices. Thus, our unexpected discovery that mitochondria are restricted to chromatin-adjacent cytoplasm as they move from the sub-spindle cytoplasm to the cortex provides a mechanism that restricts mitochondrial streaming to being perpendicular to the long axis of the spindle orientation.

How mitochondrial flow is restricted to this chromatin-associated region is not fully understood at this point. Our data suggests a model whereby MYO19 and chromatin cooperate to establish the midzone channel. This involves MYO19 tethering mitochondria to the cytoplasmic actin meshwork below the spindle preventing participation in bulk cytoplasmic streaming. Concomitantly, at the spindle midzone we propose chromatin induces distinct actin dynamics that result in MYO19-mediated active transport of mitochondria restricted to the chromatin-adjacent cytoplasm. This activity effectively draws mitochondria from the cytoplasmic actin net below the spindle to the spindle midzone and around to the cortex (see **Fig. 7f**, control).

The known role of chromatin-derived Ran-GTP on local Formin-induced actin polymerization ^45, 46^ provides support for this idea. It is also consistent with mitochondrial movement observed when MYO19 function is disrupted. In this case, uncoupling of the cytoplasmic pool of mitochondria from the actin network allows participation in cytoplasmic streaming. The cytoplasmic flow would result in passive bulk movement of mitochondria from the sub-spindle cytoplasm up and around the entire spindle as is observed in MYO19-Tail domain expressing oocytes (see **Fig. 7f**, MYO19-Tail domain expression). Thus, chromatin defines the spatial position of the channel while MYO19 enforces its exclusivity and together they ensure that mitochondrial delivery to the cortex is funneled via the chromatin-mediated corridor.

It is increasingly appreciated that the actin cytoskeleton is critical in organelle inheritance in dividing cells ^23, 47, 48, 49, 50^ and recent studies have shown that actin waves and comets in mitotic cells ensures equal and random mitochondrial inheritance ^23^. This function is not applicable in the highly asymmetric cell divisions in oocytes where the polar body is destined to degenerate ^51^. Instead, mitochondrial streaming away from the spindle in the plane opposite to that of cell division may minimize the loss of active mitochondria in the polar body at the time of cell division. This may be beneficial for subsequent embryo development which appears to be modulated by mitochondrial distribution, numbers and activity ^10, 19, 20, 52, 53, 54, 55, 56^.

Our study provides the first mechanistic details into understanding the patterning of mitochondrial distribution in mature oocytes and may provide new approaches to evaluating oocyte quality and subsequent developmental competence in the clinical setting.

## Materials and methods

### Oocyte collection and culture

All animal experiments in this study were approved by the Monash University Animal Ethics Committee and conducted in accordance with the Australian National Health and Medical Research Council (NHMRC) Guidelines on Ethics in Animal Experimentation.

3–4-week-old C57B6/J or PhAM (photo-activatable mitochondria) mice^57^ were superovulated by intraperitoneal injection of 5 IU of pregnant mare’s serum gonadotropin (Prospec) followed 44-48 hr later by intraperitoneal injection of 5 IU of human chorionic gonadotropin (hCG) (MSD Animal Health). 12-13 hr after hCG, oviductal cumulus masses were released into pre-warmed M2 medium (Sigma-Aldrich) supplemented with 300 µg/mL hyaluronidase (Sigma-Aldrich) to remove cumulus cells. Oocytes with a first polar body were transferred to drops of M2 medium under mineral oil for further experiments.

### Drug treatments

Acute drug treatment was applied to droplets of M2 medium on a confocal microscope stage to induce cytoskeletal depolymerization. Oocytes were treated with 500nM cytochalasin D (CCD; Sigma-Aldrich) or 10µM nocodazole (Sigma-Aldrich). To induce monopolar spindle formation, 5µM S-trityl-L-cysteine (STLC, Sigma-Aldrich) was added to the M2 droplet within the imaging chamber. Corresponding volumes of dimethyl sulfoxide (DMSO, Sigma-Aldrich) were used as vehicle controls.

### *In vitro* transcription and microinjection

Capped mRNA was generated by in vitro transcription of linearized plasmid templates using the T7 mMessage mMachine Transcription kit (Invitrogen) or HiScribeⓇ T7 ARCA mRNA kit (New England Biolabs) according to the manufacturer’s instructions. In selected cases, poly(A) tails were added using a Poly(A) tailing kit (Invitrogen). Transcripts were purified using the MonarchⓇ RNA cleanup kit (New England Biolabs) and stored at −80°C. mRNAs injected were: *mtPA-GFP* (500 ng/µL), *Myo19-WT-mCherry* (500 ng/µL), *Myo19-Tail-mCherry* (500 ng/µL), *Ran-WT-mRFP1* (300 ng/µL, Addgene 59750), *Ran-T24N-mRFP1* (300 ng/µL, Addgene 104561), *GFP-UtrCH* (1 µg/µL, Addgene 26737).

Oocytes were microinjected with mRNA using an electrophysiology-based picopump (PV820, World Precision Instruments) and a micromanipulator (MMN-1, Narishige)^58^. Following microinjection, oocytes were incubated in M2 medium under mineral oil (RT, 10 min) before being transferred to a heat block (+ 37°C, 10 min) to facilitate oocyte retrieval. Oocytes were cultured in M2 medium for at least 3 hr for transgene expression.

### Spindle manipulation

Oocytes were pre-incubated in culture medium containing 2 µg/mL CCD (+ 37°C, 30 min) before being transferred to M2 medium supplemented with 2 µg/mL CCD, 7% polyvinylpyrrolidone (PVP) (Sigma-Aldrich) or inactivated Sendai virus, HVJ-E (CF001EX, Ishihara Sangyo Kaisha) on a 50 mm glass-bottom dish (MatTek). All manipulations were performed on a warmed stage (Okolab) at 37°C using a Nikon Ti-2 inverted microscope equipped with Narishige micromanipulators, a dynamic biopsy laser (Octax, Vitrolife), and a spindle visualization system (Oosight, Hamilton Thorne).

Oocytes were positioned with a holding pipette adjacent to the polar body, ensuring the spindle was visible within the cytoplasm near the polar body. A small hole was drilled in the zona pellucida opposite the polar body. An enucleation pipette was inserted through this opening, aspirating the spindle into the pipette. For enucleation experiments, the pipette was then withdrawn from the cytoplasm, allowing the oocyte membrane to seal behind it.

For spindle translocation or transplantation, we adopted a previously described protocol^59^. The extracted karyoplast, held within the enucleation pipette, was exposed to the HVJ-E drop for approximately 10 sec. The pipette was then inserted into the zona through the pre-drilled opening, and the karyoplast was deposited into a position opposite its original location. Fusion occurs within 15 min.

### Live cell imaging

#### Mitochondria

To visualize mitochondria, oocytes were incubated in M2 medium containing 25 nM Tetramethylrhodamine, Methyl Ester, Perchlorate (TMRM, Invitrogen) (+ 37°C, 30 min). Oocytes were then transferred to M2 drops containing a lower concentration of TMRM (5 nM) for live-cell imaging within the imaging chamber. Imaging was performed using a Leica TCS SP8 confocal microscope equipped with either a 20x dry, 40x water immersion, or 63x oil immersion objective, or a Zeiss 980 LSM with Airyscan 2 using a 40x or 63x water immersion objective at 37 °C

#### Tubulin

To visualize meiotic spindles, oocytes were incubated in M2 drops supplemented with either 300 nM SiR-Tubulin (CY-SC002, Cytoskeleton) or 200 nM LIVE Red-Tubulin (Abberior) (+37°C, 30 min).

#### mtPA-GFP

Oocytes expressing mitochondrial photoactivatable (mtPA)-GFP were imaged approximately 3 hr after microinjection. ROIs near the meiotic spindle were exposed to brief pulses of a 405 nm laser, followed by time-series imaging using a 488 nm laser.

#### Immunofluorescence

Oocytes were fixed then permeabilized in 4% paraformaldehyde (Sigma-Aldrich) containing 0.5% Triton X-100 (Sigma-Aldrich) (RT, 20 min). Oocytes were then blocked for 1 hr in 3% bovine serum albumin in phosphate-buffered saline (PBS) before immunostaining. Oocyte were stained with: Alexa Fluor 488-conjugated mouse anti-α-Tubulin (322588; Invitrogen) or Phalloidin Alexa Fluor 555 (A34055, Invitrogen). Oocyte were then washed three times in PBS containing 0.01% Triton X-100 and 0.1% Tween 20, then transferred to PBS-PVP droplets on a glass-bottomed fluorodish (World Precision Instruments) for imaging. Confocal imaging was then performed (SP8, Leica).

### Analysis

#### Quantification of mitochondrial streaming

##### Velocity-Field Estimations of Mitochondrial Streaming

Live-cell time-lapse image sequences of mitochondria were converted to dense (i.e. per-pixel) velocity fields using computer-vision–based optical flow analysis. Inter-frame motion was estimated with Farnebäck’s dense optical-flow algorithm using MATLAB’s Computer Vision Toolbox (functions opticalFlowFarneback with estimateFlow; R2020a or later). For flow estimation, each frame was intensity-normalized, background-flattened, and then a light denoising step was applied to improve conditioning of the motion fit. The Farnebäck estimator was used to return a dense 2-D velocity vector at every pixel, which was then converted to a velocity field in µm/min using the microscope calibration (0.188 µm/pixel) and the acquisition frame interval (30 sec/frame). We interpreted the horizontal and vertical velocity components as Cartesian velocities, V_x_ and V_y_, respectively. The calculated velocity field was then used to quantify (i) the time-averaged velocity components ⟨Vx⟩ and ⟨Vy⟩; (ii) the magnitude of the time-averaged vector |⟨v⟩| = sqrt(⟨Vx⟩² + ⟨Vy⟩²), which captures persistent directional flow; and (iii) flow streamlines. It is noteworthy that the performance of this image-based kinematic analysis can be affected by occlusions, rapid intensity changes, or severe blur in the frames. The multiscale (pyramidal) polynomial modeling^60^ and temporal aggregation used here help mitigate these effects while preserving biologically meaningful transport patterns, in line with robustness considerations in classical gradient-based optical flow formulation^61, 62^. All results are reported in µm/min with a fixed Cartesian convention (+x rightward, +y upward).

##### Kymograph analysis

Time-lapse images were acquired as previously described. Line ROIs were selected from the spindle or the non-spindle hemisphere. The line selection in the spindle hemisphere traversed the meiotic spindle and the thin cortical mitochondrial layer. Kymograph images of mitochondria streaming were generated using the default kymograph function in FIJI.

##### Quantification of white pixels

To assess mitochondrial displacement over time with inhibitors, TMRM-labeled oocytes were imaged every 3 min for 3 hr and pre-processed as previously described. Oocyte images taken at 0, 1, and 2 hr after the initiation of imaging were selected for analysis.

Each image was pseudo-colored to represent different time points (red, green, or blue), with overlapping regions appearing as white pixels upon merging. The pixel distribution was quantified in Python normalized to a percentage, and analyzed using the following script:

from PIL import Image
import numpy as np
im = Image.open(’.png’)
na = np.array(im)
colours, counts = np.unique(na.reshape(−1,3), axis=0, return_counts=1)
print(colours)
print(counts)

#### Quantification of mitochondrial population in the spindle and non-spindle hemispheres

To assess mitochondrial distribution, TMRM-labeled oocytes were imaged at 3 min intervals for 3 hr, and images from the final time point were selected for analysis. The oocyte images were divided into 12 sections (30-degree intervals) using the Radial Profile Angle plugin developed in FIJI. TMRM intensity was measured in each section, normalized so that the total across all sections equaled one, and visualized using a polar histogram plot generated in R.

#### Mitochondrial membrane potential

Oocytes from PhAM mice were labeled with 5 nM TMRM. TMRM signals were excited using 561 nm and collected using a 588- to 605-nm bandpass. Simultaneously, mito-Dendra2 signals were excited at a 488-nm laser and collected from 496 to 540 nm. Z-stacks were acquired at 1.5 µm intervals across the oocytes. Oocytes oriented with the spindle in the largest cross-sectional region were selected, and maximum projection was generated using 5 images that spanned the spindle. A Gaussian filter was applied to remove noise and saturated pixels. Ratiometric analysis (TMRM/mito-Dendra2) was performed using FIJI following previously established protocols^28^, and the ratio profile was measured along a 30-pt-wide line spanning from the spindle across the oocytes.

As a control to confirm that any observed ratio differences were due to MMP rather optical or expression artifacts, oocytes were injected with mRNA encoding 4xmts-mScarlet ^63^ alongside mito-Dendra2. The same imaging and ratiometric analysis were applied.

#### Numerical simulation of fluid dynamics

To test how cell geometry and actin-driven forces shape cytoplasmic streaming, we built a simple 3D numerical model of a mouse oocyte with a cortical spindle using COMSOL Multiphysics V6.0. The oocyte was represented as a 75 µm diameter sphere, matching our experimental observations, and the spindle as a rigid ellipsoid (major axis 25 µm and minor axis 12 µm) positioned adjacent to the cortex in the “spindle hemisphere” with its long axis parallel to the cortex. Using a low Reynolds number model, we treated the cytoplasm as a viscous (dynamic viscosity µ ≈ 10^2^ Pa.s), incompressible fluid in the creeping-flow regime ^32^. All surfaces were modelled as rigid boundaries. In all simulations, the flow was driven by prescribed forces/slip velocities chosen to match the measured mitochondrial streaming velocities (or their order of magnitude), as quantified using the dense optical flow analysis. We simulated three scenarios:

##### Case A

To evaluate whether geometry alone can generate the observed flow field, a localized volumetric force was applied in the cytoplasm directly beneath the spindle and directed towards the overlying cortex (**Fig. 6a,d**). This represents an upward “pushing” of cytoplasm in the spindle hemisphere without imposing any preferred direction along the cortex. Both cortex and spindle were no-slip.

##### Case B

To mimic a cortical actin cap that uniformly drives tangential flow around the spindle, we imposed a slip velocity along the cortical cap overlying the spindle, directed tangentially in all azimuthal directions (“360°” around the spindle), while the rest of the cortex remained no-slip (**Fig. 6b,e**). The slip-velocity profile and angular extent of the active cap were chosen so that cortical velocities in the spindle hemisphere matched our experimental end-on flow maps, and the location of the slip domain followed the tangential cortical flow versus spherical polar angle reported by Liao and Lauga ^32^.

##### Case C

To mimic a cortical actin cap that selectively drives tangential flow along the spindle minor axis, we imposed anisotropic tangential slip over the cortical cap, with slip strongest perpendicular to the spindle major axis and reduced along it (**Fig. 6c,f**). This represents an anisotropic actin network whose activity is aligned relative to the spindle. The cortex outside the cap and the spindle surface were no-slip.

For each case, we computed the steady-state flow field and then extracted end-on and side-on sections matching our imaging planes. We also solved a transient advection–diffusion equation for a passive tracer concentration field to model mitochondrial redistribution driven by the computed flow field, with tracer initially confined to a sub-spindle volume. The tracer diffusivity was set to D ≈ 0.017 µm^2^/sec, based on diffusion coefficients reported for mitochondria in large vertebrate embryos^64^. Velocity vectors and tracer concentration maps were then compared with the experimental data to identify the forcing scenario that best reproduced the observed streaming patterns.

### Cytoplasm PIV analysis

To quantify and visualize oocyte flow magnitude and direction, Particle Image Velocimetry (PIV) analysis was performed using the open-source MATLAB software PIVlab (version 3.12.001) along with custom written MATLAB scripts. Data preparation involved cropping individual oocytes and generating a pixel-specific mask to restrict analysis to the oocyte boundaries. The brightfield channel served as the primary PIV input. The PIV process utilized a two-pass, decreasing interrogation area approach. The size for the initial pass was calculated as approximately five times the inverse of the pixel size, measured in micrometers per pixel. Pass 2 reduced the interrogation area to 50% of Pass 1. A 50% step size was used relative to the interrogation area implemented for both passes respectively. Vector fields were calculated using the Gauss 2×3 point sub-pixel estimation method and the ‘Extream’ correlation robustness setting. Instantaneous vector fields were subsequently temporally averaged to generate a temporal mean vector field, from which overall flow velocity magnitudes and angles were extracted.

For visualization, a custom MATLAB script incorporated a modified version of the quiverc function (https://www.mathworks.com/matlabcentral/fileexchange/3225-quiverc) to overlay color-coded vectors (by magnitude) onto the raw data. Quantification of flow velocity magnitudes was restricted to regions of interest exhibiting characteristic swirling flow patterns. For Nocodazole-treated oocytes, the SYBG channel was used to guide the definition of quantified regions of interest.

### Statistical analysis

For experiments comparing two groups statistical significance was determined using an unpaired t-test. Significance levels were represented as follows:

p <0.05 (*), p<0.01(**), p<0.001 (***), and p<0.0001 (****).

In the violin and box plots, the central line represents the median, while the bottom and top of the line correspond to the 25% and 75%, respectively.

## Acknowledgement

We thank Dr. Wai-Shan Yuen for the contributions to the initial experiments and Dr. Jeffrey Mann for the critical comments on the manuscript. We thank Dr. Minc Nicolas for insightful discussions. We also acknowledge the Monash Animal Research Platform and the Monash Micro Imaging Platform for their support in enabling the experiments performed herein. This study was funded by the ARC DP160104892 and NHMRC 1165627 and 200112.

## Author contributions

I.-W.L. and J.C. designed the research. I.-W.L., and J.C. wrote the manuscript. I.-W.L., performed the experiments. R.S. performed micromanipulation experiments and Z.-W.W, M.R.D and G.S, contributed to the early inception of the project. I.-W.L. contributed new reagents and analytic tools; I.-W.L., M.N., S.U., J.Y., S.A., and R.N. analyzed the experimental data. J.C. and D.A. acquired funding.

---

**Extended Data Fig 1.**
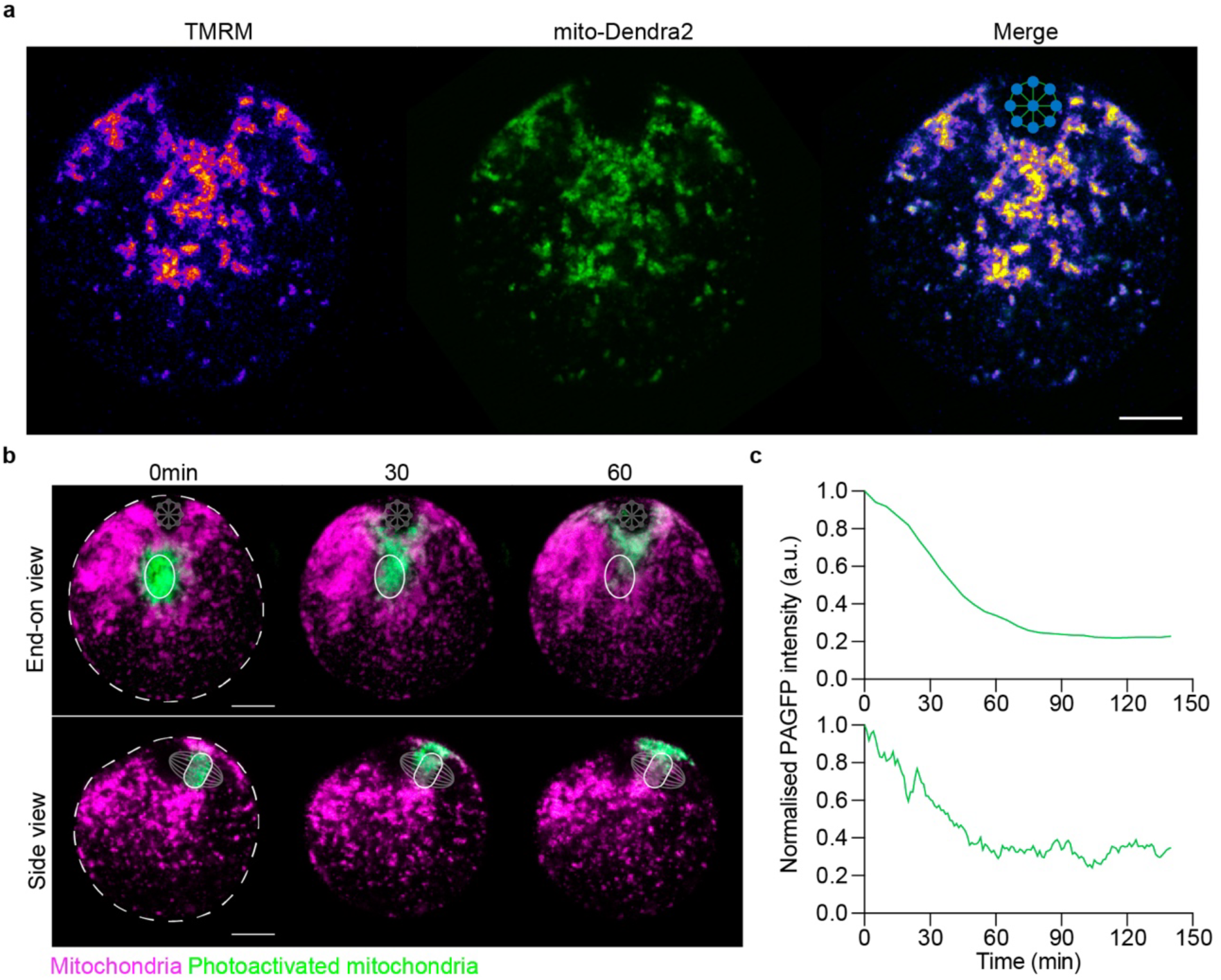
Photoactivated mitochondria within the mitochondrial mass near the meiotic spindle show continuous movement. **a,** Representative images showing mitochondrial distribution labeled with TMRM (fire LUT) and mito-Dendra2 (green) in the end-on view of the oocyte. Scale bar, 15 µm. **b,** Live-cell images of oocytes expressing mtPA-GFP shown in spindle end-on view (top) or side view (bottom). Time is indicated in min with time point 0 representing the first frame post-photoactivation. Magenta, mitochondria (TMRM); green, photoactivated mitochondria (mtPA-GFP). White circle indicates the initial photo-activated region. S, spindle; the dashed line, the oocyte membrane. Scale bar, 15 µm. Data represents 12 oocytes from three replicates. **c**, Quantification of the normalized fluorescence intensity of mtPA-GFP at the ROI where mtPA-GFP was initially photoactivated. Decrease in fluorescence over time indicates mitochondrial movement out of the activated regions.

**Extended Data Fig. 2:**
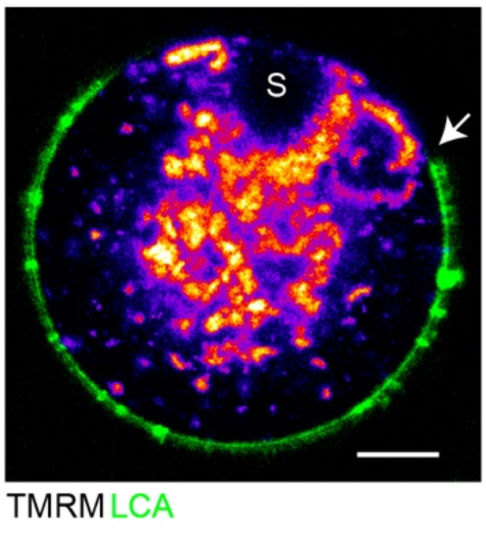
Mitochondria streaming in the cortex stops and turns inward at the boundary of the polarised cortex. Mitochondria (TMRM) and microvilli (LCA) were labelled in MII oocytes. The spindle is located at 12 o’clock, indicated by ‘S’. Arrowhead indicates the border between the polarised microvillus-free and microvillus-rich cortex. Note that cortical mitochondria are restricted to the microvillus-free cortex. Scale bar, 10um. Data represents 40 oocytes from 3 replicates.

**Extended Data Fig 3:**
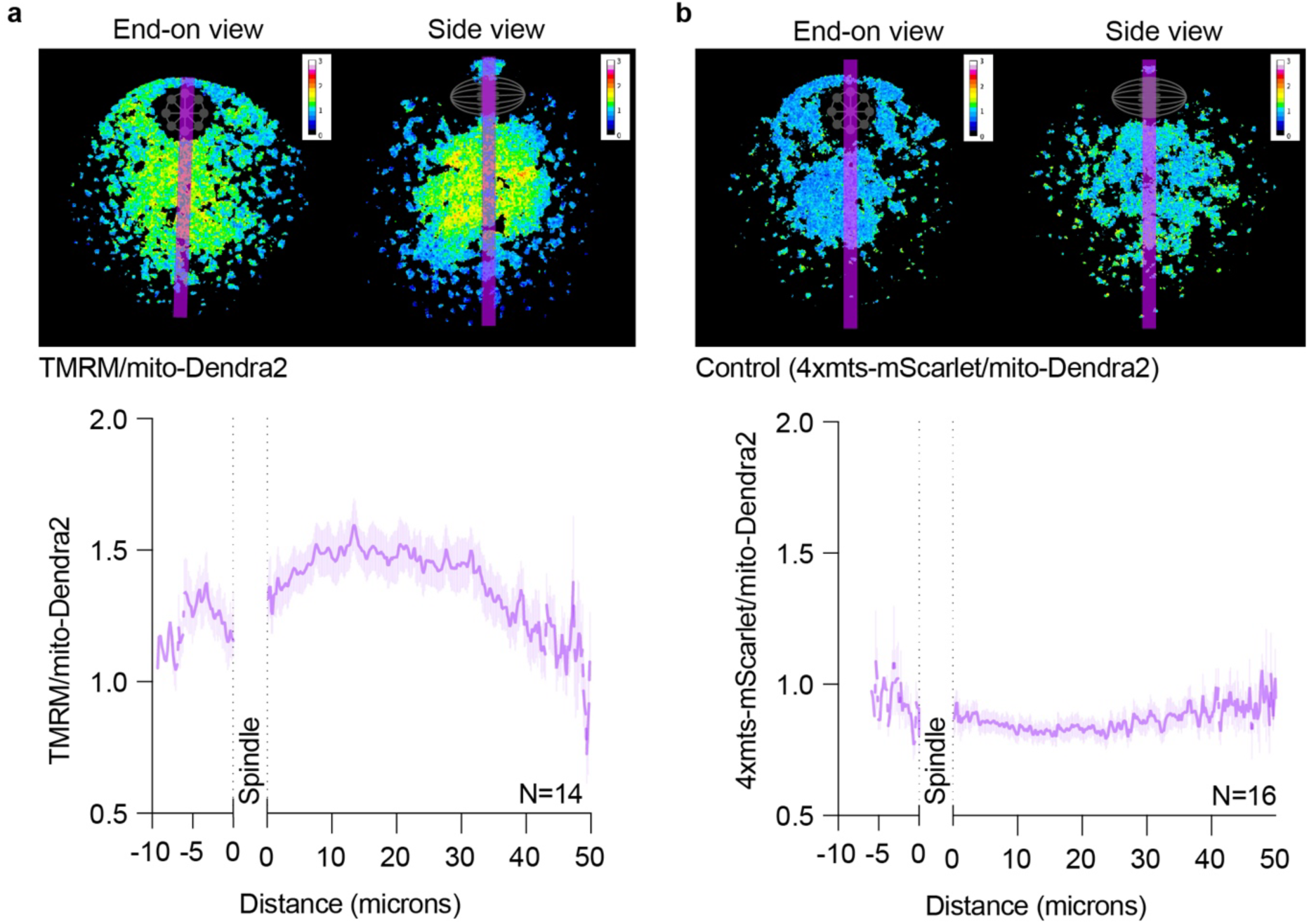
MMP forms a gradient from the spindle hemisphere to the non-spindle side of the oocyte. **a,** Representative ratiometric images of TMRM/mito-Dendra2 fluorescence intensity provide a spatial map of MMP from the spindle hemisphere to the non-spindle hemisphere. The spindle is indicated by a spindle pictogram. A pseudo-color scale displays high MMP (warm colors, high ratio) and low MMP (cool colors, low ratio). The line plot below shows the TMRM/mito-Dendra2 ratio profile measured along the 30-pixel-wide purple line shown in the image. Data are represented as mean ± SEM. **b**. Ratiometric control analysis using two MMP-independent indicators. Shown are representative ratiometric images and line plot profile of 4xmts-mScarlet/mito-Dendra2 fluorescence intensity. No gradient is observed, confirming that the MMP gradient in (**a**) is not due to imaging artifacts. Data are represented as mean ± SEM.

**Extended Data Fig 4.**
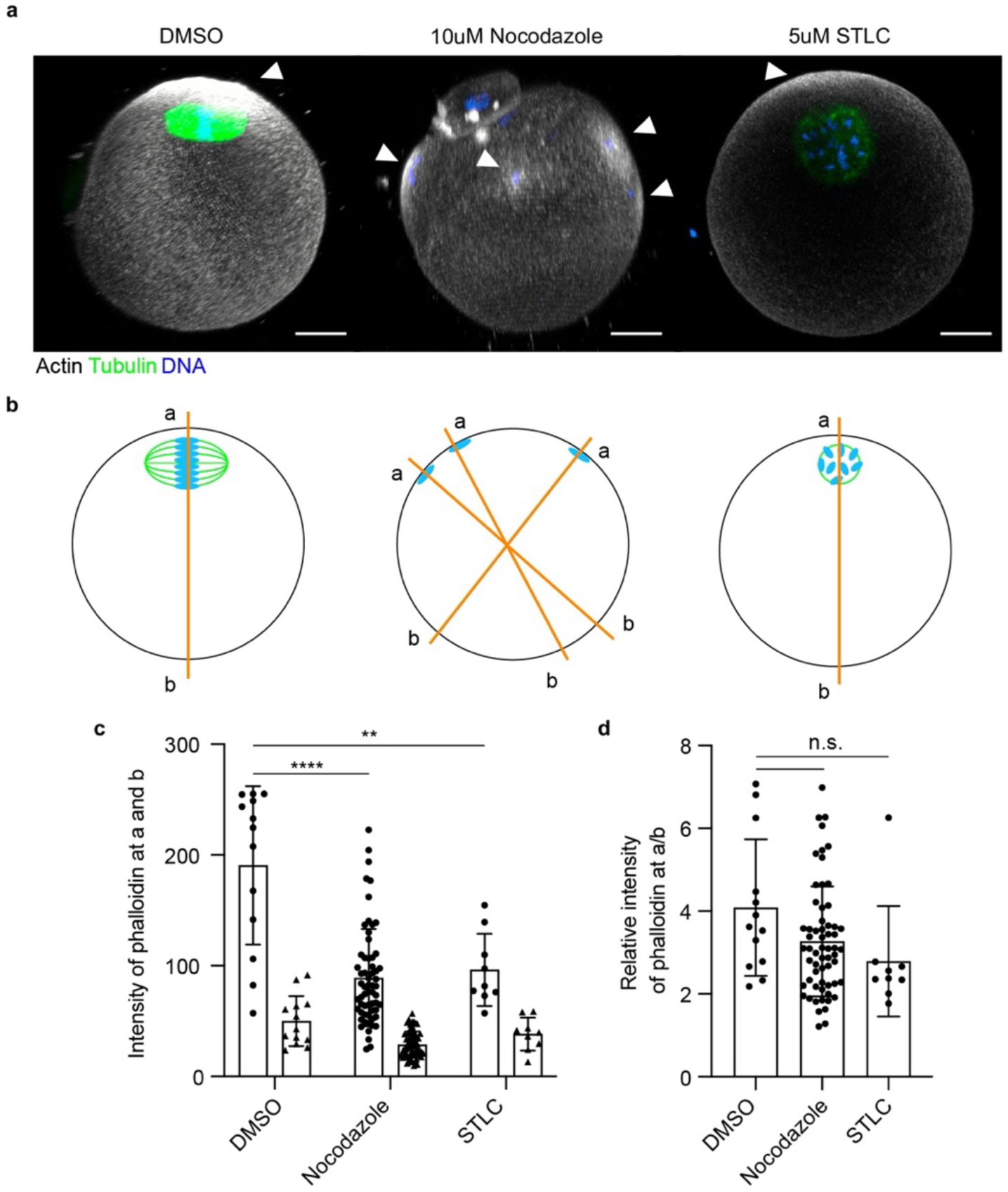
Cortical actin cap persists in oocytes treated with nocodazole or STLC. **a.** Representative images showing the distribution and intensity of cortical actin in MII oocytes after 4 hr of incubation with DMSO, nocodazole, or STLC. Images are 3D projections generated from 115 slices acquired at 1 μm intervals. In DMSO-treated oocytes, the bipolar spindle is typically positioned near the cortical actin cap. In Nocodazole-treated oocytes, spindle disruption results in chromosomes dispersed along the cortex with multiple cortical actin caps forming over individual chromosomes. In STLC-treated oocytes, a globular spindle forms and is displaced away from the cortex. Scale bar; 15 μm. **b.** Schematics showing the positions where phalloidin intensity was measured in DMSO, Nocodazole, or STLC-treated oocytes. ‘a’ indicates the cortex close to the chromosome. ‘b’ indicates the cortex non-spindle to ‘a’. **c.** Quantification of phalloidin intensity at regions ‘a’ and ‘b’. Unpaired t-test (****p<0.0001, **p<0.01). **d.** Ratio of phalloidin intensity comparing region ‘a’ to ‘b’ (a/b). Unpaired t-test (n.s., not significant).

**Extended Data Fig. 5.**
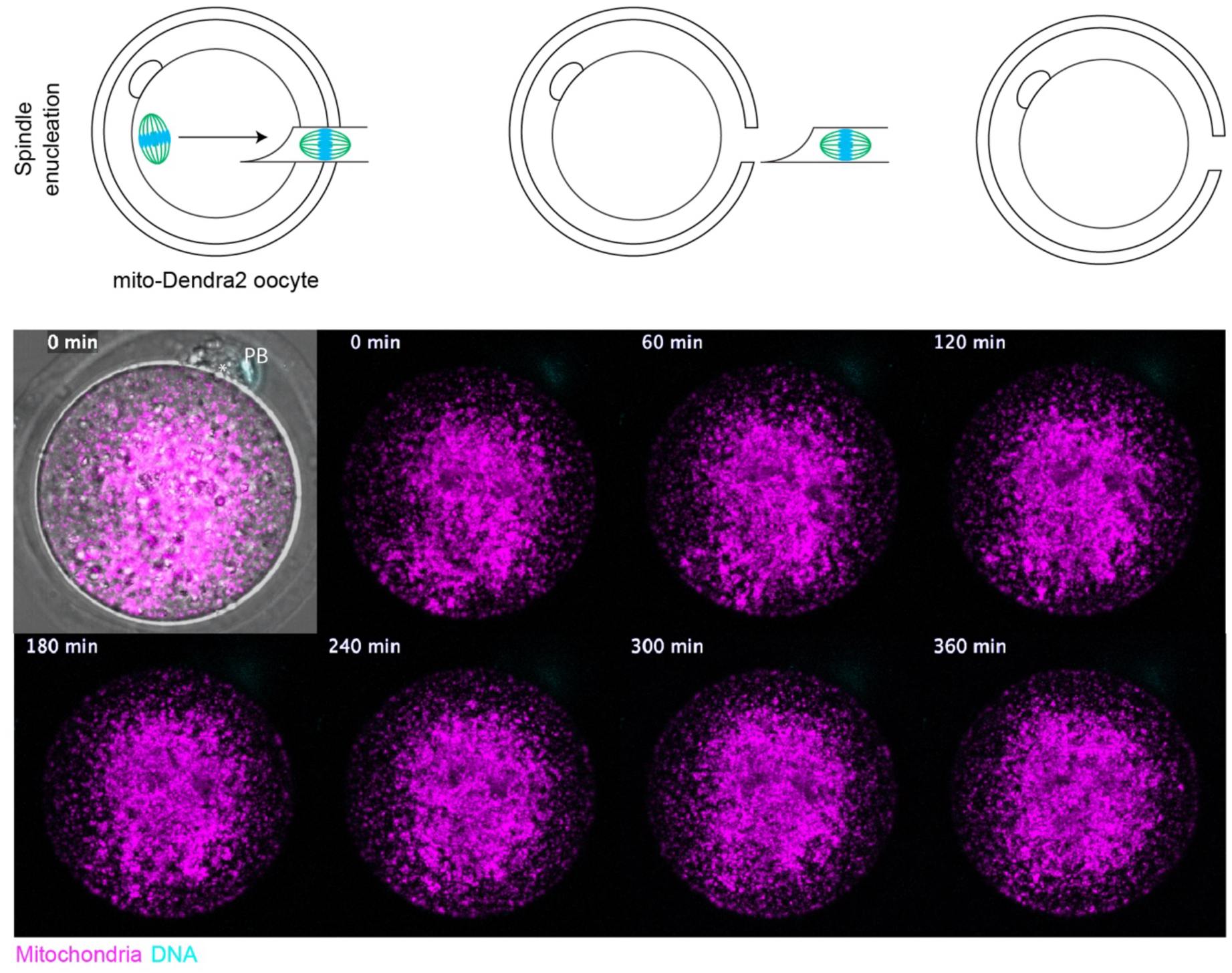
Mitochondria cease bulk movement and become less dynamic after spindle enucleation. Time series of mitochondrial distribution in an enucleated oocyte. Images are maximum projections of 8 z-slices acquired at 1.5 μm intervals. Mitochondria (visualized with mito-Dendra2) exhibit no dynamic movement in the ooplasm, and the streaming pattern is not observed. DNA is labeled with the Abberior DNA live dye.

**Extended Data Fig. 6:**
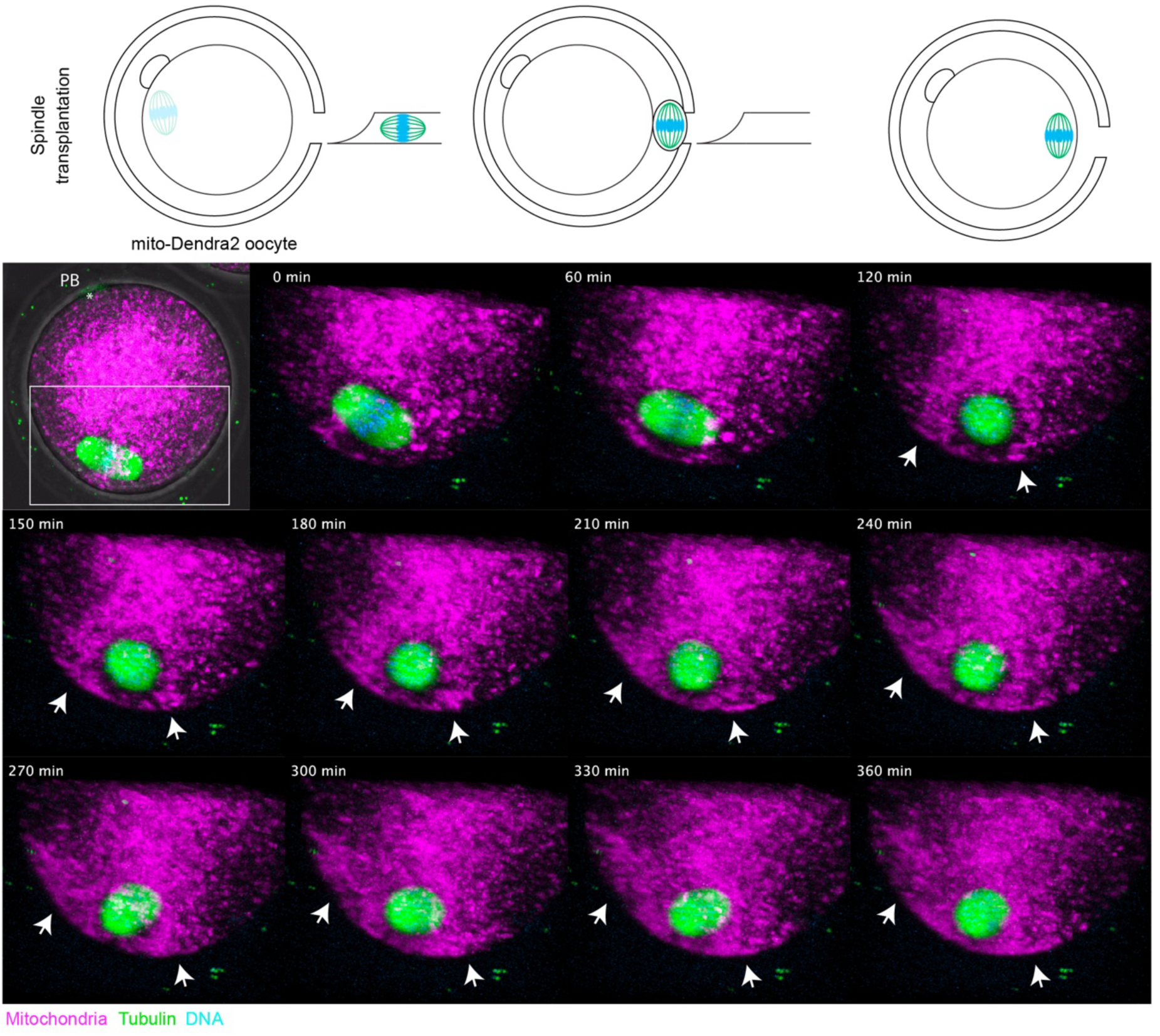
Mitochondria establishes bulk movement following spindle translocation. Time series of mitochondrial distribution in a spindle-translocated oocyte. Images are 3D projections of 20 z-slices acquired at 1.5 μm intervals. Mitochondria (visualized with mito-Dendra2) exhibit bulk movement towards the spindle (indicated by white arrows). Tubulin is labeled with SiR-Tubulin; DNA is labeled with the Abberior DNA live dye.

**Extended Data Fig. 7.**
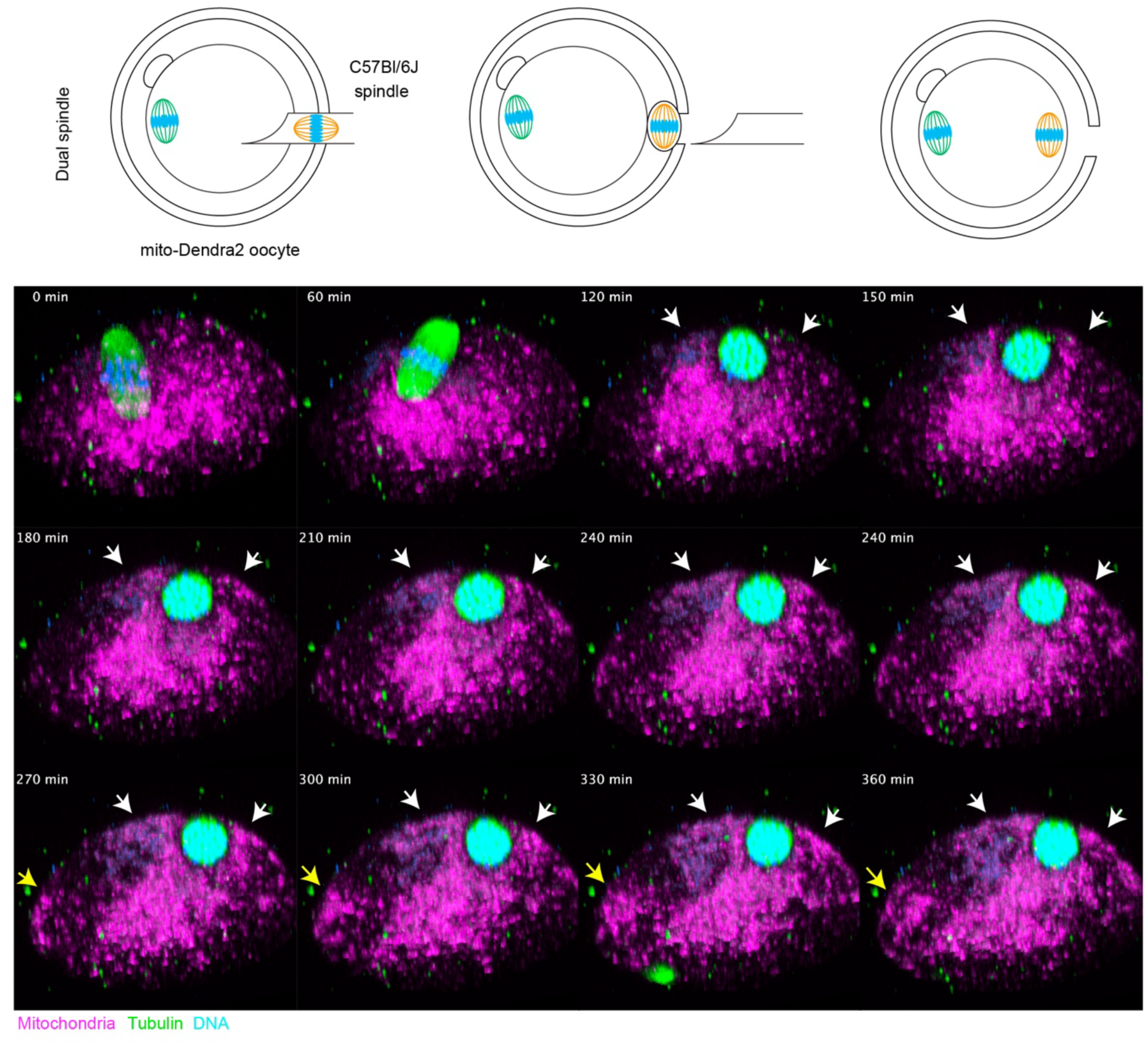
Mitochondria establish bulk movement near the transplanted spindle. Time series of mitochondrial distribution in a dual-spindle oocyte. Images are 3D projections of 20 z-slices acquired at 1.5 μm intervals. Mitochondria (visualized with mito-Dendra2) exhibit bulk movement towards the donor spindle (indicated by white arrows). Partial mitochondrial streaming driven by the original recipient spindle is also observed (indicated by yellow arrows). Tubulin is labeled with SiR-Tubulin; DNA is labeled with the Abberior DNA live dye.

**Extended Data Fig 8:**
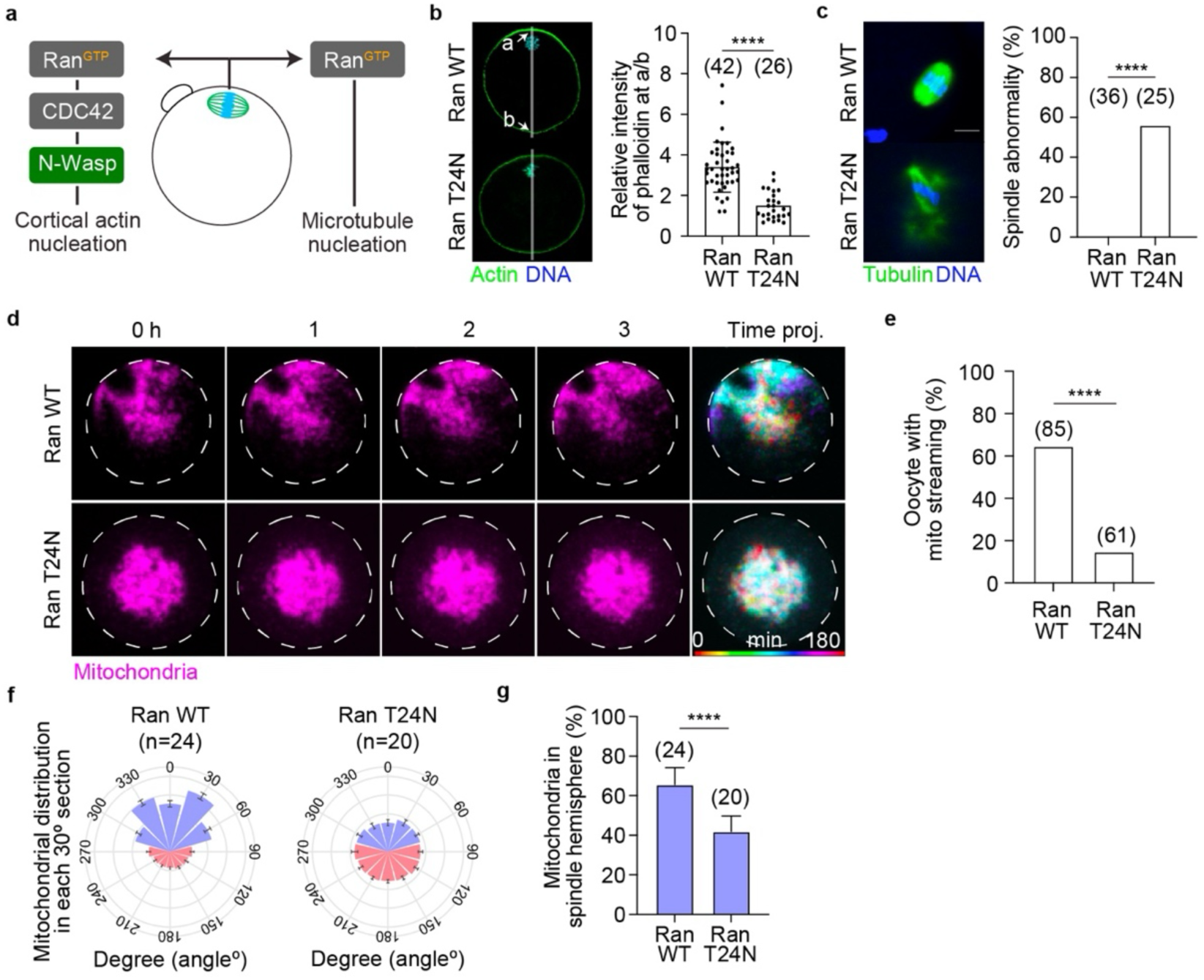
Mitochondrial streaming is regulated by the activity of Ran-GTP emanating from chromosomes. **a,** Schematic illustration showing the role of Ran-GTP in forming the cortical actin layer and microtubules in MII oocytes. **b,** Representative images of cortical actin and quantification of phalloidin intensity in Ran WT- or Ran T24N-expressing oocytes. Green, actin (phalloidin); blue, chromosome (Hoechst). ‘a’ refers to measurements in the spindle cortex, while ‘b’ indicates the non-spindle cortex. Unpaired t-test (****p<0.0001). **c,** Representative images of spindle and quantification of abnormal spindle formation after Ran T24N expression. Chi-square test (****p<0.0001). **d,** Representative live-cell and time-projection images of mitochondrial distribution (TMRM) in oocytes expressing Ran WT or T24N. Dashed line indicates the oocyte membrane. **e,** Percentage of oocytes with mitochondrial streaming patterns after Ran WT or T24N expression. Chi-square test (****p<0.0001). **f,** Quantification of the mitochondrial population measured in each hemisphere (as described in Fig. 3f). Blue represents mitochondrial distribution in the spindle hemisphere (150 degrees), while red indicates those on the non-spindle side (210 degrees). **g,** Percentage of mitochondria localized in the spindle hemisphere relative to total mitochondrial population measured in **f**. Data are expressed as the mean ± SD. Unpaired t-test (****p<0.0001).

**Extended Data Fig. 9:**
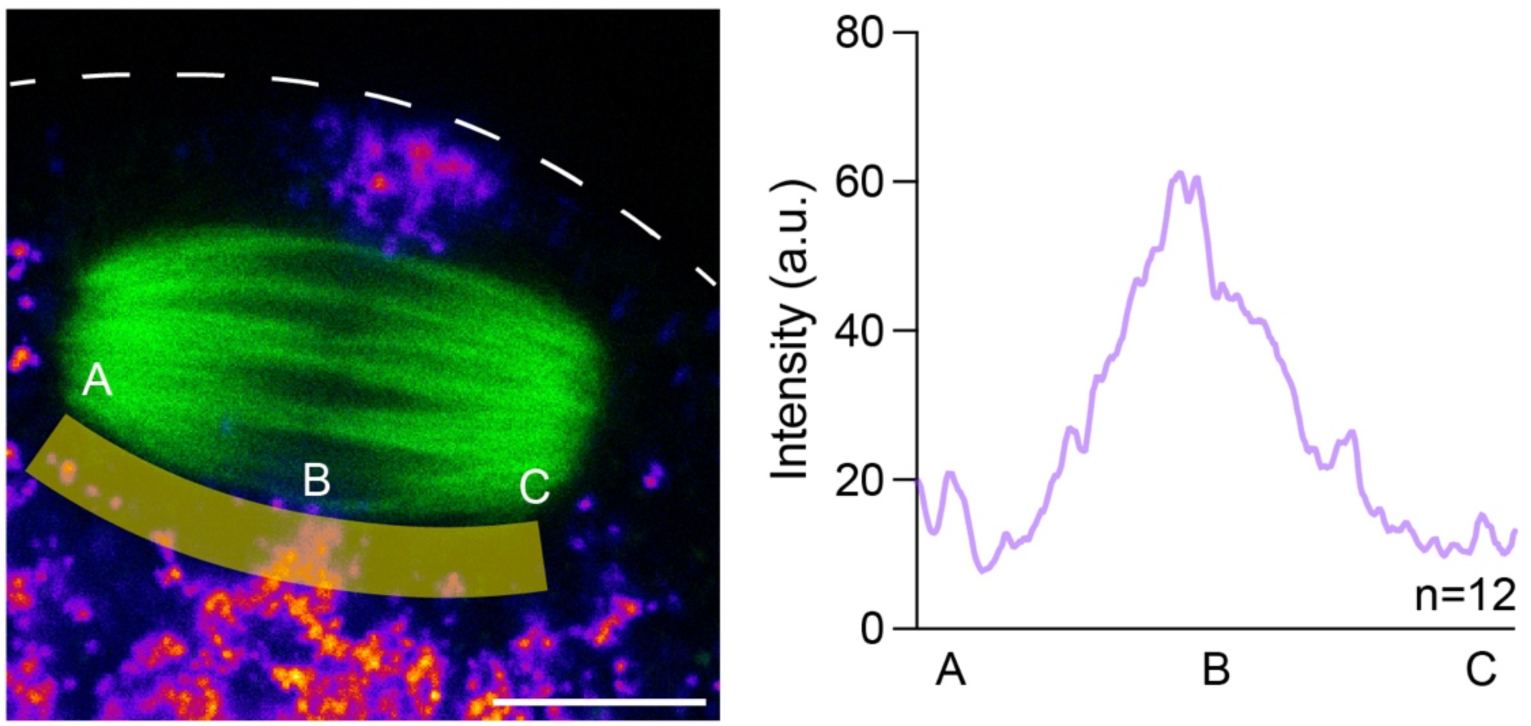
Mitochondrial distribution along the immediate sub-spindle regions is restricted to the midzone. Representative image of the spindle (green, tubulin) and mitochondria (fire, mito-Dendra2). The yellow shaded regions indicate the area used for fluorescence intensity profiling along the length of the spindle below from the pole A through the midzone B to pole C. The dashed white line indicates the cortex. Scale bar: 10 μm. Right panel shows the quantification of mito-Dendra2 intensity along this region.

**Extended Data Fig. 10:**
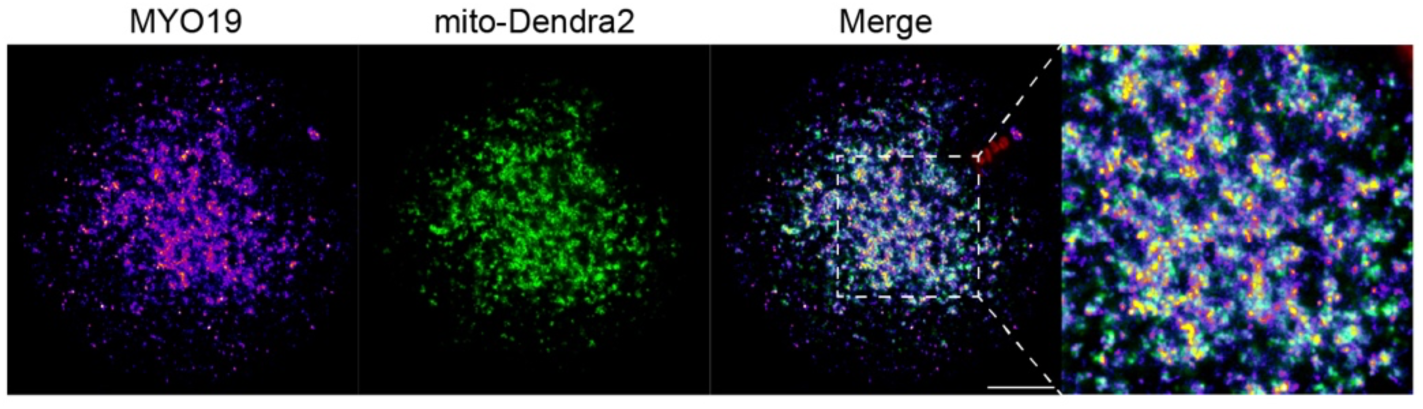
MYO19 localizes with mitochondria in MII oocytes. Representative immunofluorescence images of an MII oocyte showing MYO19 (fire), mito-Dendra2 (green), DNA (red). MYO19 localizes with mito-Dendra2-labelled mitochondria in the sub-spindle cytoplasm. The dashed box indicates the region shown a higher magnification demonstrating colocalization of MYO19 puncta with mitochondria. Scale bar: 15 μm.

**Extended Data Fig. 11:**
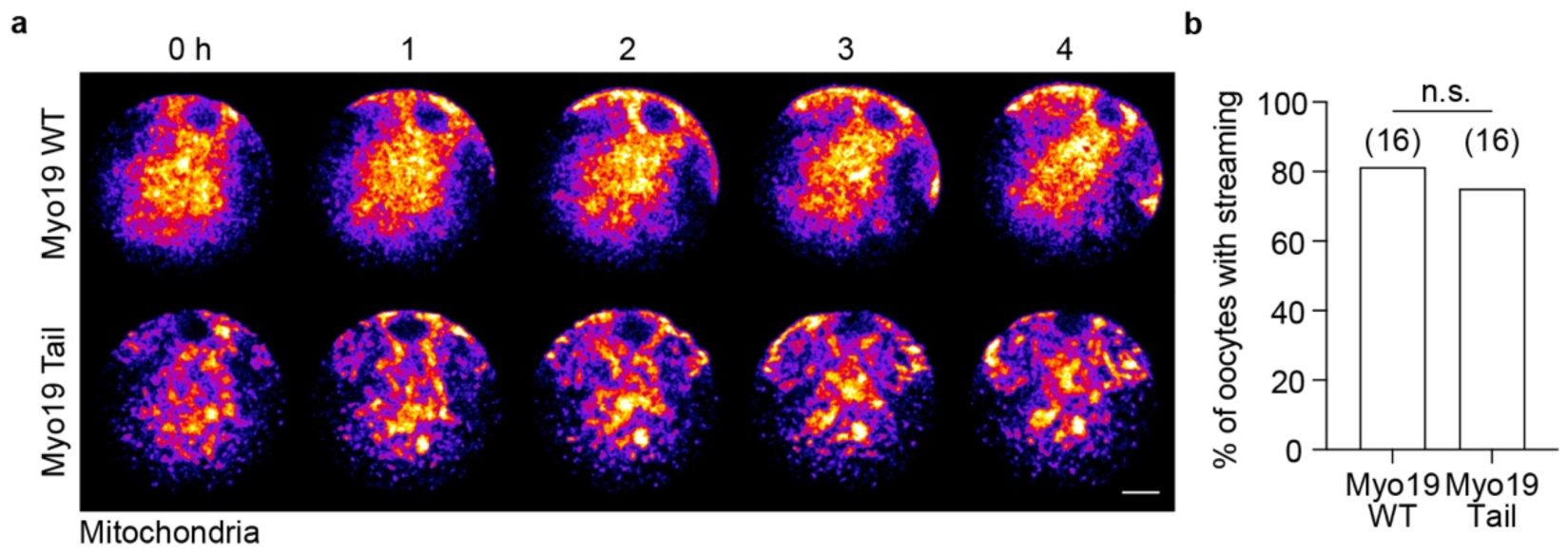
Mitochondrial streaming persists in the presence of *Myo19*-Tail expression. **a,** Representative time-lapse images of mitochondrial streaming (mito-Dendra2) in oocytes expressing *Myo19*-WT or *Myo19*-Tail. Data are representative of 16 oocytes per group. Scale bar, 15 µm. **b,** Percentage of oocytes displaying mitochondrial streaming patterns in *Myo19*-WT or *Myo19*-Tail. Chi-square test (n.s.: non-significant).

**Supplementary Video 1. Spindle orientation-dependent patterns of mitochondrial streaming at the spindle hemisphere of the MII oocyte.** Mitochondria showed less directional movement at the spindle hemisphere in the side view spindle oocyte, while they dynamically moved toward the top of the spindle by traversing either side of the spindle and sliding down along the cortex in the end-on view oocyte. Images were captured over a period of 3 hr (Δt = 30 sec). Mitochondria were labeled with TMRM. Scale bar, 15 µm.

**Supplementary Video 2. Mitochondria move toward isolated chromosomes in the presence of nocodazole.** Mitochondria labeled with TMRM (fire LUT) and chromosomes (green) with SYBR Green. Three representative movies are shown. Images were captured at Δt = 30 sec.

**Supplementary Video 3. Mitochondria streaming in the DMSO-treated oocyte.** 3D projection movie of an oocyte labeled with TMRM (magenta) and SiR-Tubulin (green) in the end-on view.

**Supplementary Video 4. Mitochondria streaming in the STLC-treated oocyte.** 3D projection movie of an oocyte labeled with TMRM (magenta) and SiR-Tubulin (green). Note the monopolar spindle configuration following STLC treatment.

**Supplementary Video 5. Mitochondria cease bulk movement and become less dynamic after spindle enucleation.** Time-lapse movie of mitochondrial distribution in an enucleated oocyte. Images are maximum projections of 8 z-slices acquired at 1.5 μm intervals. Mitochondria (visualized with mito-Dendra2) exhibit no dynamic movement in the ooplasm, and the streaming pattern is not observed. DNA is labeled with the Abberior DNA live dye.

**Supplementary Video 6. Mitochondrial streaming is re-established after spindle translocation.** Time-lapse movie of mitochondrial distribution in a spindle-translocated oocyte. Images are 3D projections of 20 z-slices acquired at 1.5 μm intervals. Mitochondria (visualized with mito-Dendra2) exhibit bulk movement towards the spindle (indicated by white arrows). Tubulin is labeled with SiR-Tubulin; DNA is labeled with the Abberior DNA live dye.

**Supplementary Video 7. Mitochondrial streaming is established near a transplanted spindle in a dual-spindle oocyte.** Time-lapse movie of mitochondrial distribution in a dual-spindle oocyte. Images are 3D projections of 20 z-slices acquired at 1.5 μm intervals. Mitochondria (visualized with mito-Dendra2) exhibit bulk movement towards the donor spindle (indicated by white arrows). Partial mitochondrial streaming driven by the original recipient spindle is also observed (indicated by yellow arrows). Tubulin is labeled with SiR-Tubulin; DNA is labeled with the Abberior DNA live dye.

**Supplementary Video 8. Mitochondrial channeling is seen in all oocytes displaying mitochondrial streaming.** 3D reconstruction of mitochondria and mitochondrial channeling as demonstrated in a typical group of oocytes. Of the 48/54 oocytes showing streaming, all exhibit the chromosome-associated channeling. The oocytes represented in this movie were collected based on the presence of a visible polar body and were randomly placed in the imaging chamber. This clearly demonstrates that oocytes display different patterns of mitochondrial streaming depending on spindle orientation. Mitochondria are labeled with mito-Dendra2 (magenta) and the spindle with SiR-Tubulin (green). Z-stacks were acquired at 1 μm intervals. Data are representative of 54 oocytes from 4 replicate experiments.

**Supplementary Video 9. The directionality of mitochondrial streaming segments the ooplasm of the spindle hemisphere.** A series of z-slices showing mitochondria labeled with TMRM (fire LUT), spindle with SiR-Tubulin (green), and chromosomes with Abberior DNA dye (blue). Note the mitochondria-rich regions perpendicular to the spindle axis and mitochondria-free regions parallel to the spindle pole.

**Supplementary Video 10. Mitochondria tightly align with spindle actin at the spindle midzone.** A series of z-slices showing mitochondria labeled with TMRM (magenta), spindle with SiR-Tubulin (green), and actin with GFP-UtrCH (gray).

